# β1 Integrin–FAK–Piezo1 signalling axis drives *in-situ* stiffening mediated ECM remodelling and invasion of 3D breast epithelium

**DOI:** 10.1101/2025.01.07.631639

**Authors:** Kabilan Sakthivel, Anna Kotowska, Ellen Juel Portner, Catherine Merry, Pontus Nordenfelt, Adam Cohen Simonsen, Amanda J. Wright, Vinay Swaminathan

## Abstract

Stiffening of tissue is a hallmark of cancer progression, driving invasive phenotypes through complex interactions between cells and their extracellular matrix (ECM). However, the mechanisms linking mechanical cues to ECM remodelling and invasion remain incompletely understood. Here, using an *in-situ* stiffening model that allows for modulation of ECM stiffness around fully formed normal mammary acini embedded in their native ECM microenvironment, we identify critical steps in basement membrane (BM) and stromal ECM remodelling during invasion and discover the molecular mechanisms driving this process. We find that stiffening of the ECM around normal mammary acini results in rapid loss and degradation of laminin (LN) and upregulation of the fibronectin (FN) secretion around the acini. This priming phase is followed by the onset of invasion which requires localized upregulation of LN production and ECM remodelling. Mechanistically, ECM production and remodelling as well as invasion is mediated by β1 integrin–FAK signalling, which activates mechanosensitive ion channels (MSCs). Further, we identify Piezo1 as the MSC downstream of β1 integrin–FAK that drives BM disruption and stromal ECM remodelling. Taken together, our results identify the mechanisms by which stiffness can trigger invasive phenotypes from normal tissues.

## Introduction

The extracellular matrix (ECM), which comprises both, the basement membrane (BM) and the stromal ECM, plays a pivotal role in tissue morphogenesis and homeostasis, influencing cellular behaviors such as proliferation, differentiation and migration [1], [2]. In the mammary gland, the mechanical properties of the ECM, such as its stiffness, are tightly regulated to drive morphogenesis and preserve tissue integrity [3]. However, during tumorigenesis, the ECM undergoes extensive remodeling, leading to changes in its stiffness and composition and compromised structural integrity [3], [4]. These biomechanical changes drive malignant phenotypes in mammary epithelial cells (MECs) through loss of tissue polarity, disruption of the BM, increased proliferation and the acquisition of migratory phenotypes [5], [6]. While considerable progress has been made in understanding how ECM stiffness promotes invasiveness in MECs and mammary acini isolated from their native ECM [7]–[10], the effects of stiffening on fully developed mammary acini embedded in its self-established ECM just as in the body, which more accurately represents tumor progression, remains poorly understood.

Normal mammary acini, composed of a polarized epithelial monolayer surrounded by a BM, serve as a robust model for studying tissue-specific responses to mechanical cues. The interplay between ECM mechanics, stromal ECM and BM remodeling, and cellular invasion is both dynamic and complex [6]. For example, a stiff microenvironment can promote MECs to remodel BM through enhanced protease activity and generation of forces on the BM resulting in its disruption [11] or through increased deposition of ECM proteins such as fibronectin (FN) [12], [13], both leading to increased invasion and tumor progression. Similarly, changes in expression of BM modifying proteins such as netrin-4 can lead to increases in its stiffness, facilitating cancer cell invasion and metastasis [14]. Cancer cells can also utilize non-proteolytic pathways to generate forces and squeeze through the pores of the ECM [11], [15], [16]. However, the dynamics of ECM remodelling and invasion processes, especially in the context of fully formed acini and the stiffening ECM surrounding it, are still not well characterized.

Mechanotransduction, the process by which cells sense and respond to mechanical stimuli, plays a central role in mediating cellular responses to ECM stiffening and integrins are key activators and transducers of cell-ECM interactions in mammary acini [17]. Integrins interact with both stromal ECM (FN, collagen) and BM (laminin (LN)) components to remodel ECM and regulate BM integrity through signaling pathways involving focal adhesion kinase (FAK) and others, several of which have been implicated in promoting tumor progression [18]–[24]. In parallel, recent studies have also highlighted the role of mechanosensitive ion channels (MSCs), including Piezo1 and transient receptor potential (TRP) channels, in cellular responses to changes in ECM stiffness, implicating them in both normal physiology and cancer progression [25]–[28]. Intriguingly, emerging evidence suggests a potential interplay between integrin-mediated signaling and MSCs in regulating stiffness-induced tumor progression [29]–[33]. However, the exact nature of this crosstalk and the specific role they play during tumour progression, both in the context of breast cancer as well as other cancers, still requires better understanding.

To answer these questions in physiological settings, here we utilized an *in-situ* stiffening model of 3D mammary acini in their native ECM environment to simulate the increase in ECM stiffness observed during tumor progression. Using an interpenetrating network (IPN) hydrogel system composed of alginate and BM extract (BME), we grew fully mature mammary acini under soft (normal) conditions and induced stiffening through *in-situ* crosslinking of the alginate once the acini were fully developed and had established their own ECM environment. This model allowed us to capture the progressive morphological changes, ECM remodeling, and mechanotransduction pathways activated during *in-situ* stiffening. Using this system, we first identified distinct phases of invasion: an initial priming phase involving BM (LN) degradation and stromal ECM (FN) upregulation lasting a few days, followed by an invasion phase marked by re-secretion of LN and reorganization of FN which triggered the onset of proliferation and migration. Moreover, using a combination of agonists, inhibitors and function blocking antibodies, we find that activation of Piezo1 is a critical downstream effector of β1 integrin-FAK signaling which is sufficient to trigger *in-situ* stiffening-induced invasion and ECM remodeling. Taken together, these results provide new mechanistic insights into the interplay between ECM dynamics, integrin signaling, and its crosstalk with MSC activation in the context of ECM stiffening during tumor progression and highlight potential therapeutic targets to mitigate the invasive transition in breast cancer.

## Results

### *In-situ* stiffening of normal mammary acini results in altered ECM composition, remodeling and distinct invasive phenotypes

To capture the process of stiffening induced changes in tissue organization, we adapted the IPN hydrogel system composed of alginate and BME to first grow 3D acini from single MECs in a mechanically soft environment [9] and then cross-linked the hydrogel *in-situ* to stiffen it after formation of a fully mature acini. We first confirmed that our system achieved the desired stiffness range through atomic force microscopy (AFM) measurements. *In-situ* crosslinking of the soft gel with 20 mM CaCl_2_, hereafter referred as the *in-situ* stiffened gel, increased stiffness from 86 ± 29 Pa to 24.8 ± 22.1 kPa (Figure 1A), effectively spanning the stiffness range of both normal and cancerous breast tissues [34]. Additionally, we prepared stiff IPN gels by premixing 20 mM CaSO_4_.2H_2_O with the IPN gel mixture [9], which resulted in a stiffness of 3 ± 4.1 kPa (Figure 1A). Next, we encapsulated single-cell suspensions of MCF10A MECs within the soft IPN gels and cultured them for 2 weeks until they formed fully mature mammary acini structures (Figure 1B). After the 2-week period, we crosslinked, and *in-situ* stiffened a subset of gels with the embedded acini by adding 20mM CaCl_2_ and allowed the acini to continue growing in unstiffened (soft) and *in-situ* stiffened gels for another 2 weeks (Figure 1B). Subsequently, all gels were fixed and stained for the nucleus (DAPI), F-actin, LN (laminin-332) and FN to measure acini morphology, BM integrity and stromal ECM deposition, respectively. In parallel, we also embedded single MECs in stiff gels and allowed them to grow for 28 days for comparison with the soft and *in-situ* stiffened gels (Figure 1B).

**Figure 1:**
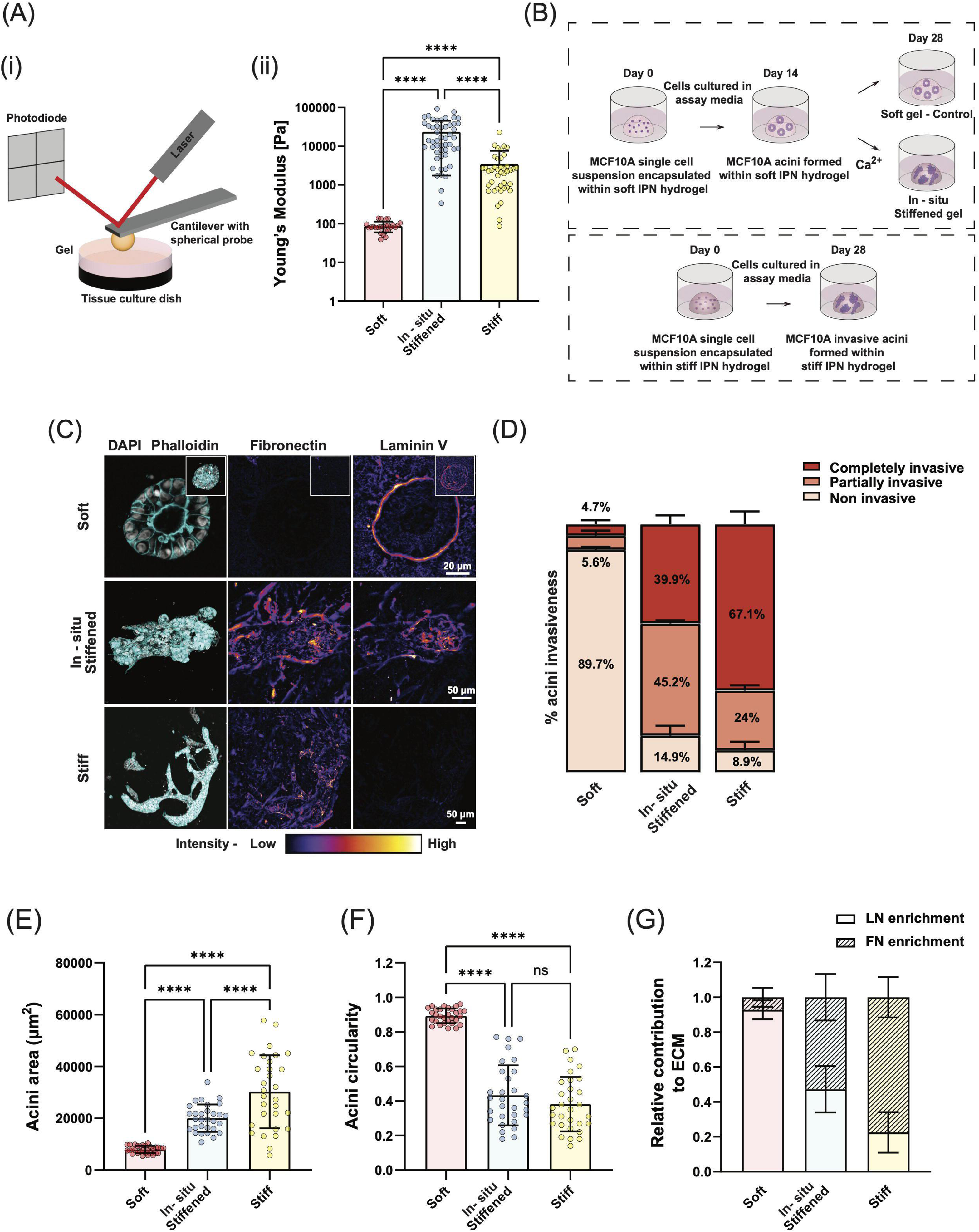
In-situ stiffening of the normal mammary acini microenvironment induces distinct invasive phenotypes and dynamic ECM remodeling. (A) Schematic representation of the atomic force microscopy (AFM) setup utilized for measuring the Young’s modulus of soft, in-situ stiffened, and stiff IPN gels. The AFM probe indents the gel surface to assess its mechanical properties across different stiffness conditions. AFM measurements of the Young’s modulus of IPN gels under different stiffness conditions (n = 2 different experiments per condition, with 3 × 3 - point measurements performed on three distinct regions in each experiment). Stiffness was measured immediately after the formation of soft and stiff gels and post-in-situ stiffening of the soft gels. (B) Schematic overview of the experimental overflow for 3D acini culture. Single-cell suspensions of MCF10A were encapsulated in soft and stiff IPN gels. Once normal mature acini formed in soft IPN gels, a subset of the soft gels was in-situ stiffened with 20 mM CaCl_2_ on day 14. Acini in unstiffened (soft control) and in-situ stiffened IPN gels were cultured for an additional 14 days. Parallelly, acini within stiff IPN gels were cultured for 28 days. (C) Confocal images showing nuclei (gray), F-actin (cyan), and FN and LN staining (fire LUT applied to highlight FN and LN levels within the acini) of acini in soft, stiffened and stiff IPN gels. Z-stacked images of acini cultured in in-situ stiffened and stiff IPN gels are shown, while a single Z-plane (middle plane of the stack) image of acini cultured in soft IPN gels is displayed to highlight lumen formation (Z-stacked image for the same acini is shown in the inset). Scale bars: 20 µm for soft; 50 µm for in-situ stiffened and stiff. (D)-(G) Quantification of (D) percentage of invasive acini, (E) acini area, (F) acini circularity, and (G) relative FN and LN enrichment in acini in soft, in-situ stiffened, and stiff IPN gels (n = 30 acini from 3 different experiments, with 10 acini per experiment for acini area, circularity, and FN and LN enrichment quantifications; n ≥ 30 acini from 3 experiments for invasiveness quantification; Ordinary one-way ANOVA test, ****p < 0.0001, ns=non-significant).

Confocal microscopy of fixed acini revealed that most mature acini after 4 weeks of culturing in the soft IPN gels remained intact and spherical with fully developed lumens at their centers (Figure 1C and Figure S1). In contrast, for *in-situ* stiffened and stiff gels, we observed most acini to be significantly larger, disorganized and without an intact lumen though we did observe some variability in the levels of response (Figure 1C, Figure S1). When we examined the deposition and remodelling of the BM and the ECM by imaging LN and FN respectively, we found that normal mature acini in soft IPN gels had a clear and distinct layer of LN encircling each acinus with relatively faint or no FN staining around it (Figure 1C). However, while both *in-situ* stiffened and stiff acini showed elevated levels as well as fibrillogenesis of FN, there were dramatic differences in LN levels between the two conditions (Figure 1C). Invasive acini in *in-situ* stiffened conditions showed elevated levels of LN staining with significantly remodelled LN that was in a discontinuous layer (Figure 1C). In contrast, in stiff gels, as previously reported [9], we found very low levels of LN along the invading fronts or anywhere else within the epithelial structure (Figure 1C). To ensure that calcium ions used to *in-situ* crosslink IPN gels did not influence the structure and production of the ECM or changes in morphology in *in-situ* stiffened gels, we added 20 mM CaCl_2_ to acini formed within BME-only gels without alginate (BME-only gels) and found that it had no effect on acini morphology or on FN and LN levels (Figure S2).

To quantify the different acini structures observed, as well as capture the heterogeneity in acini response, we classified each acini as either non-invasive (completely spherical, with an intact lumen), partially invasive (loss of spherical integrity to some extent, filling up of lumen/loss of lumen and invading cells sprouting out of the actual acini area) and completely invasive (complete loss of spherical integrity)(Fig 1F, Figures S1, S3). This analysis revealed that while normal mammary acini grown in soft IPN gels were ∼90% non-invasive, stiffening of gels around fully formed normal acini resulted in ∼85% of the embedded acini to undergo morphological transformation with varied levels of invasion into the surrounding gel (Figures 1D, 1E, 1F and Figure S1). Specifically, ∼40% lost their spherical structure, had filled-in lumens and were completely invasive, while the rest exhibited partial invasiveness, characterized by protrusions sprouting from their surfaces while still maintaining their circular morphology and lumens (Figures 1D, 1E, 1F and Figure S1). In contrast, growing single MECs in stiff IPN gels resulted in ∼90% of the acini to be invasive with a majority of these, ∼67%, being completely invasive with loss of lumen and no remaining spherical morphology (Figures 1D, 1E 1F). Within *in-situ* stiffened gels, the deposition and remodeling of FN and LN varied according to the acini’s response to the stiffening process (Figure S4). Non-invasive acini exhibited robust LN signal encircling their periphery, indicative of an intact BM, and displayed minimal FN deposition. In partially invasive acini, FN levels were elevated, particularly at the invasive fronts, accompanied by a weak and discontinuous LN signal around the periphery, reflecting BM distortion (Figure S3). Fully invasive acini, however, exhibited high levels of both FN and LN deposition, with these ECM components prominently localized at the periphery and within the acini (Figure S3). To quantify these differences in ECM composition, we measured the overall background-subtracted intensities of LN and FN staining and plotted the relative contribution of each component to the ECM in the acini i.e their relative levels within the ECM (Figure 1G, Figure S4). This analysis showed that while normal mammary acini were ∼80-90% enriched with LN, *in-situ* stiffening of the ECM around mature acini resulted in nearly equal contribution levels of the two (∼50%) which was mediated by partial reduction in LN deposition and significant increase in FN secretion (Figure 1G, Figure S4). In contrast, growing single MECs in stiff gels resulted in very low levels of LN leading to an ECM ∼80% enriched with FN (Figure 1G, Figure S4). To examine whether the observed response of normal MECs to *in-situ* stiffening is also applicable to tumorigenic cell lines, we repeated the above experiments with tumorigenic MECs - MCF10A DCIS.com cell line. Consistent with previous reports, MCF10A DCIS.com acini, while maintaining a circular morphology in soft gels, failed to form lumens and displayed multiple breaches in the BM (Figure S5) [35], [36]. Most importantly, similar to normal MCF10A acini, MCF10A DCIS.com acini responded to *in-situ* stiffening by significantly increasing its area, reducing its circularity and invading into the gel (Fig S5). Concurrent with these changes, *in-situ* stiffening also resulted in changes in the ECM composition surrounding the invading acini with similar levels of FN and LN enrichment (Fig S5).

Together, these findings demonstrate that *in-situ* stiffening of normal mammary acini within their native BM-ECM induces invasive phenotypes through significant changes in production and remodelling of this native ECM, a process that is distinct from those of acini grown in pre-stiffened environments.

### *In-situ* ECM stiffening results in distinct phases of changes in ECM composition, remodeling and onset of invasion in mammary acini

Having established a system that allows for *in-situ* stiffening of fully mature normal acini in their native ECM environment, we next aimed to investigate the timeline of morphological and ECM remodeling events during this process. To do so, we first mapped the dynamics of stiffening of the IPN gel using particle tracking microrheology (PTM) [37], [38]. Briefly, 6 μm diameter polystyrene beads were embedded in soft IPN gels, incubated for 24 hours, and then imaged on a widefield microscope with a high acquisition rate, ∼300 Hz, for several minutes to observe the Brownian motion and diffusion of the beads. After collecting data on the soft gels over 22 hours (measurements taken every 2 hours for 10 hours per day), the gels were crosslinked using 20 mM CaCl_2_ to induce stiffening and the embedded beads were further tracked over 2 days. For each set of tracked data, the mean square displacement (MSD) of the bead over time was determined and used to measure the low frequency plateau of the elastic modulus G’_0_ (Fig 2A, see methods). Plotting G’₀ over time revealed that the gels underwent a pronounced increase in stiffness within the first 6 hours (∼3.5-fold increase compared to the soft gels), followed by a continued increase in stiffness up to 24 hours post-crosslinking, after which the stiffness plateaued and remained constant for the subsequent 24 hours (Fig 2A). These measurements show that *in-situ* crosslinked IPN gels undergo rapid stiffening within the first 6 hours and then remain stable over timescales lasting days.

**Figure 2:**
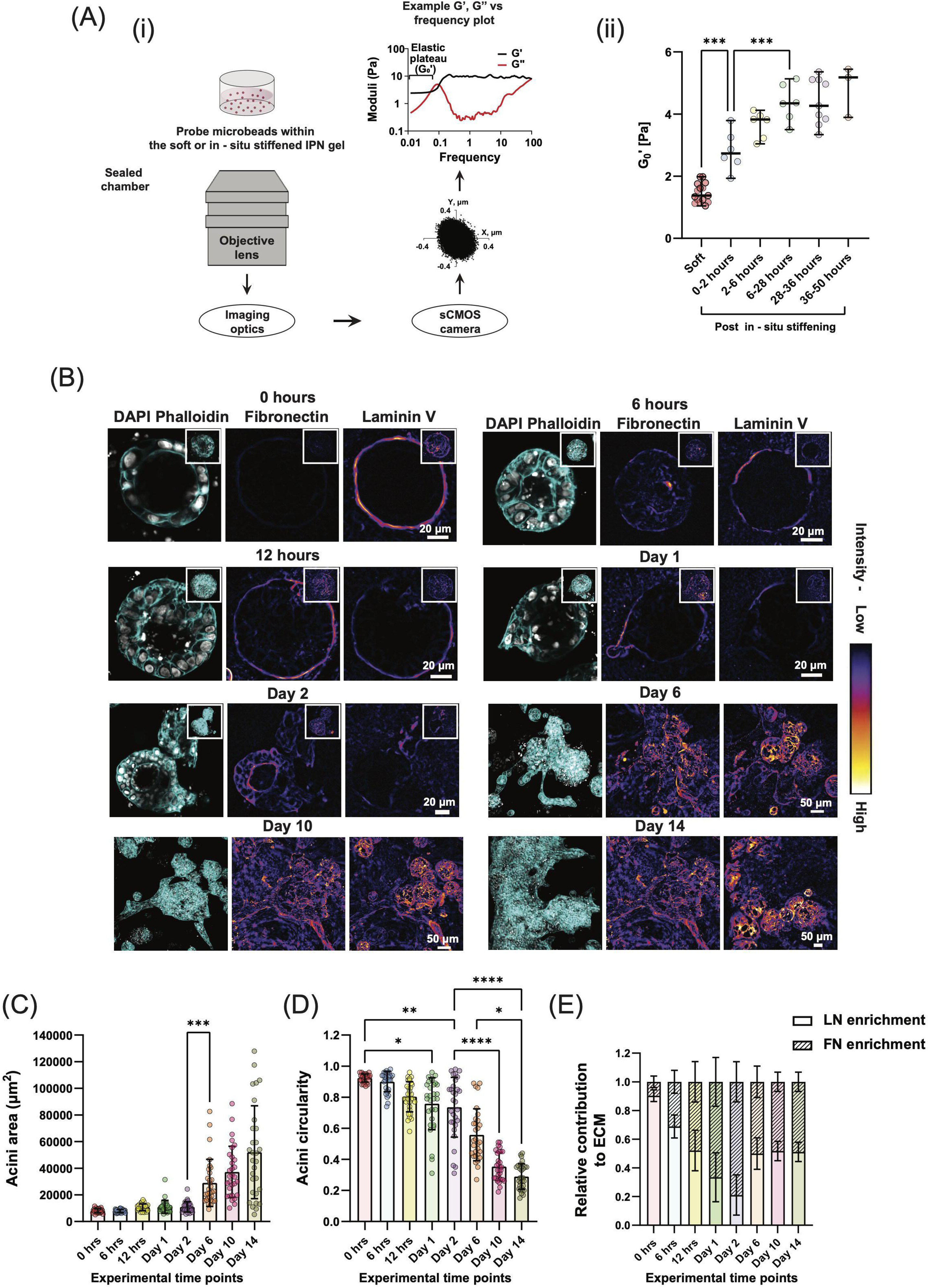
In-situ ECM stiffening results in distinct phases of changes in ECM composition, remodeling and onset of invasion in mammary acini. (A) (i) Schematic representation of the imaging setup used for PTM to measure the local stiffness of soft, IPN gels before and after the in-situ stiffening process. The schematic includes an example of a small region of interest image of the probe, where the Brownian motion of the probe is tracked using a center-of-mass algorithm to extract the low-frequency elastic modulus (G’₀) of the material surrounding the probe. (ii) Low-frequency elastic modulus (G’₀) of the soft IPN gel measured via PTM before and after the in-situ stiffening process (n = 2 gels per condition, with 3 beads in different regions of each gel; Ordinary one-way ANOVA test, ***p < 0.001). Stiffness measurements for the soft IPN gels were recorded over the first 24 hours post-gelation. The same gels were subsequently in-situ stiffened, and measurements were taken over the following 2 days to monitor changes in stiffness post-modulation. (B) Confocal images of nuclei (gray), F-actin (cyan), and FN and LN (fire LUT applied to highlight FN and LN levels within the acini) staining at various time points post-in-situ stiffening of soft IPN gel. Z-stacked images of acini within in-situ stiffened IPN gels are shown for Days 6, 10, and 14 post-stiffening. Single Z-plane images (middle plane of the stack) are displayed for time points 0 hours, 6 hours, 12 hours, Days 1, and 2 post-stiffening to illustrate lumen filling and early invasion events (Z-stacked images for the same acini are shown in the insets). Scale bars: 20 µm for time points - 0 hours, 6 hours, 12 hours, Days 1 and 2, 50 µm for time points - Days 6, 10, and 14, 50 µm. (C)-(E) Quantification of (C) acini area, (D) acini circularity, and (E) relative FN and LN enrichment in acini at various time points post-in-situ stiffening of the soft IPN gel (n = 30 acini from 3 different experiments, with 10 acini per experiment for acini area, circularity, and FN and LN enrichment quantifications; n ≥ 30 acini from 3 experiments for invasiveness quantification; Kruskal-Wallis test, *p < 0.05, **p < 0.01, ***p < 0.001, ****p < 0.0001).

Next, we embedded single MCF10A MECs in soft IPN gels and allowed them to form normal mature acini for 14 days just as above. After *in-situ* crosslinking with 20mM CaCl_2_, subsets of *in-situ* stiffened gels were fixed at 6 hours, 12 hours, 24 hours, 2 days, 6 days, 10 days and 14 days post stiffening and stained for DAPI, F-actin, FN and LN. Confocal imaging and quantification of acini morphology and LN and FN levels under these conditions revealed a remarkably fast process of changes in ECM levels and a slower process of changes in acini morphology and invasion upon stiffening (Figures 2B, 2C, 2D, 2E and Figure S6). At 6 hours post-stiffening, we observed a robust increase in FN levels around the periphery of the acini with concurrent reduction in LN levels, which reduced the enrichment of LN levels by ∼2 fold (Figures 2B, 2E and Figure S6). The acini however remained intact and circular during this initial 6-hour period with no increase in area or loss of circularity (Figures 2C. 2D). At 12 hours post-stiffening, we started to observe luminal infiltration in some acini and while there was some indication of an increase in its area and loss in circularity, this wasn’t significant, however the LN levels continued to significantly reduce, and FN levels continued to increase in the surrounding ECM (Figures 2B, 2C, 2D, 2E and Figure S6). This process continued in samples fixed 24 hours and 48 hours post-stiffening, and by 48 hours, we observed cells beginning to invade into their surrounding environment with high FN signal localized around and through the invading front and no LN signal at these regions (Figures 2B, 2C, 2D, 2E and Figure S6). Quantification revealed only a small change in acini area and circularity at 48 hours with most of the LN gone from around the acini resulting in a FN enriched ECM (with an enrichment index of ∼0.8) (Figures 2B, 2C, 2D, 2E and Figure S6).

This first phase was followed by a second phase observed in samples fixed 6 days post-stiffening, where we started to observe changes in the pattern. LN levels now seemed significantly higher compared to 2 days which was confirmed by quantification resulting in the acini being equally enriched for LN and FN levels (Figure 2B and Figure S6). This enrichment was primarily driven by upregulation of LN levels while FN levels remained high and stable (Figure S6). The acini themselves lost most of their circular morphology and had invaded significantly into their surrounding with high FN and LN levels at the invading fronts (Figures 2B, 2C, 2D, 2E and Figure S6). The acini continued to invade and expand, were no longer circular and high levels of both, FN and LN were maintained during subsequent timepoints until day 14 (Figures 2B, 2C, 2D, 2E and Figure S6).

Taken together, these results suggest that stiffening of normal mature mammary acini first results in rapid changes in stromal ECM and BM levels, where stromal ECM production increases, and BM level is downregulated or degraded. During this phase, the acini remain mostly intact and spherical with some cellular infiltration into the lumen. This is then followed by a second phase which is characterized by onset of significant invasion which coincides with upregulation of BM levels and significant remodelling of both, the BM and the stromal ECM.

### β1 Integrin–FAK axis regulates ECM composition and remodeling during *in-situ* stiffening-induced acini invasion

Mechanical signals from the ECM are transduced into biochemical responses primarily through integrins and their downstream effectors, such as FAK, which play critical roles in cell-matrix interactions and mechanotransduction. Prior studies have highlighted the role of integrin-mediated signaling in driving invasive phenotypes in MECs [19]–[21], [39]. Given our observations of dramatic ECM remodeling, including LN disruption and FN fibrillogenesis in *in-situ* stiffened acini, we hypothesized that integrin-FAK signaling might mediate stiffening-induced morphological changes and invasive transitions. To test this, we investigated the expression and activation of β4 integrins [40], which primarily bind LN, and β1 integrins which interact with FN [39], in acini grown in soft, *in-situ* stiffened, and stiff IPN gels.

To do so, we first stained acini grown in soft, *in-situ* stiffened and stiff IPN gels for total β4 integrins and active β1 integrins that bind to FN. Confocal imaging of normal acini in soft conditions showed a distinct circular and continuous localization of β4 integrins surrounding the intact acini that resembled LN staining in Figure 1 (Figure 3A). Even though β1 integrins can dimerize with α6 and α3 integrins to bind to LN [41], staining for the β1 subunit in soft gels showed a faint and relative to β4, insignificant signal around the acini as well as everywhere else in proximity of the epithelial structure (Figure 3A). This was confirmed through quantification of intensity and levels of active β1 integrins (relative to total β4) which showed overall low levels of β1 activation (Figures 3B, 3C, 3D, Figures S7A, S7B, S7C). In comparison to the acini in soft gels, we found unexpected changes in integrin levels in acini within stiff and *in-situ* stiffened gels (Figures 3A). Firstly, both *in-situ* stiffened and stiff acini with invasive phenotypes continued to exhibit elevated levels of total β4 integrins (Figure 3A), which contrasted with the observed differences in LN levels between acini in stiff and *in-situ* stiffened gels. In addition, not surprisingly, high levels of active β1 integrin staining was now observed throughout the invasive acinar regions correlating with the FN localization pattern in Figure 1 (Figure 3A). This led to an overall significant increase in enrichment of active β1 integrins at the acini-ECM interface in *in-situ* stiffened and stiff conditions compared to acini grown in soft gels (Figures 3B, 3C). These results suggest that *in-situ* stiffening of normal mature acini results in significantly increased levels of active β1 integrins with no change in total β4 integrin levels.

**Figure 3:**
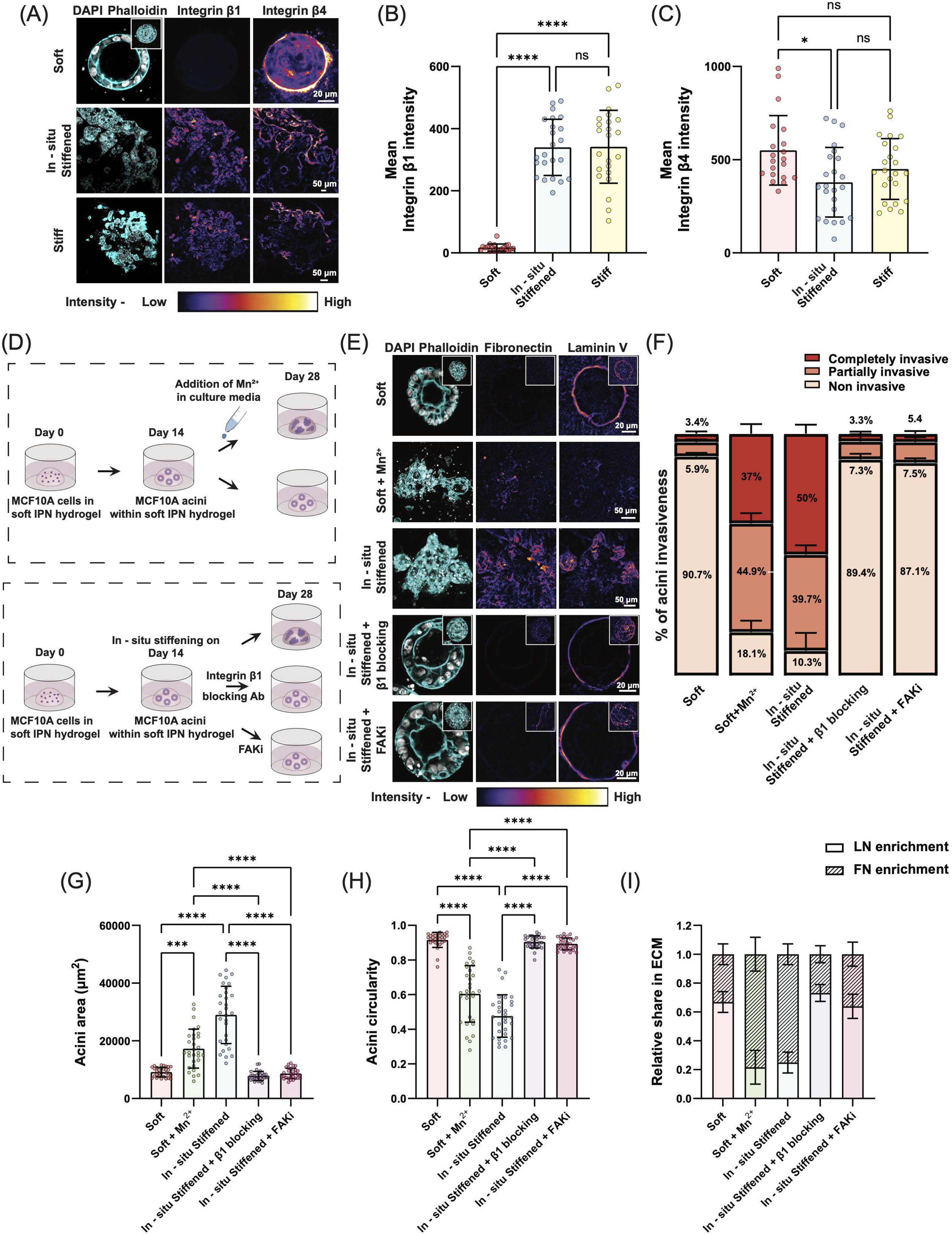
β1 Integrin–FAK Axis Drives ECM Remodeling and In-situ Stiffening-Induced Invasion. (A) Confocal images of acini cultured in soft, in-situ stiffened, and stiff IPN gels, showing nuclei (gray), F-actin (cyan), β1 integrin, and β4 integrin (fire LUT applied to highlight β1 and β4 integrin expression). Z-stacked images are displayed for in-situ stiffened and stiff gels, while a single Z-plane image of acini in soft gels highlights lumen formation (Z-stacked image for the same acini in inset). Scale bars: soft, 20 µm; in-situ stiffened and stiff, 50 µm. (B-C) Quantification of mean (B) FN and (C) LN intensities within acini cultured in soft, in-situ stiffened, and stiff IPN gels (n = 20 acini from 2 different experiments, with 10 acini per experiment; Kruskal-Wallis test, *p < 0.05, ****p < 0.0001, ns = not significant). (D) Schematic representation of the experimental workflow to investigate integrin-dependent phenotypic alterations in acini. After mature acini formation in soft IPN gels, gels were: (i) left untreated (soft control), (ii) treated with MnCl_2_ to activate integrins, (iii) in-situ stiffened and either left untreated (in-situ stiffened control), or treated with integrin β1 blocking antibody or FAKi. (E) Confocal images showing nuclei (gray), F-actin (cyan), and FN and LN (fire LUT applied) in acini cultured under various experimental conditions. Z-stacked images are shown for acini in soft + Mn²⁺ and in-situ stiffened gels, while single Z-plane images highlight lumen formation and retention in soft, in-situ stiffened + β1 blocking, and in-situ stiffened + FAKi conditions (Z-stacked image for the same acini in inset). Scale bars: 20 µm for soft, in-situ stiffened + β1 blocking, and in-situ stiffened + FAKi; 50 µm for soft + Mn²⁺ and in-situ stiffened. (F-I) Quantification of (F) percentage of acini invasiveness, (G) acini area, (H) acini circularity, and (I) relative FN and LN enrichment in different experimental conditions (n = 30 acini from 3 different experiments, with 10 acini per experiment for area and circularity quantifications; n ≥ 30 acini from 3 experiments for invasiveness; Kruskal-Wallis test, ***p < 0.001, ****p < 0.0001).

Since our data above indicated a role for activation of integrins, we next tested if activation was sufficient to result in invasive phenotypes in normal mammary acini. To test this, we treated fully mature normal acini grown in soft gels with 0.5mM MnCl_2_ that promotes the activation of integrins independent of stiffness (Figure 3D) [42]. Upon MnCl_2_ treatment, we found that ∼88% of the normal mature acini in soft gels now exhibited an invasive phenotype as quantified by significant loss of acini circularity and increase in area (Figures 3E, 3F, 3G, 3H, 3I Figures S7D, S7E). However, compared to acini in *in-situ* stiffened gels, MnCl_2_ treated acini expanded less and did not produce FN to the same level (Figures 3D, 3I, Figure S7F). Furthermore, we found that LN levels in these acini were also significantly lower compared to acini in *in-situ* stiffened gels (Figures 3D, 3I, Figure S7G). This suggests that while integrin activation is sufficient to promote invasion of mammary acini in the soft environment, *in-situ* stiffening activates other pathways or amplifies integrin pathways to promote the complete invasive phenotype.

We next investigated the specific role of β1 integrins in this process by inhibiting the activity of β1 integrins using the specific inhibitory antibody AIIB2 or its downstream target FAK during *in-situ* stiffening of normal mature acini (Figure 3D). After two weeks of culturing in the *in-situ* stiffened gel, mammary acini in the presence of the integrin activation blocking antibody or the FAK inhibitor did not undergo any invasive transformation, maintaining their confined area, circularity, lumen and FN and LN levels similar to those in the soft control group (Figures 3E, 3F, 3G, 3H, 3I, Figures S7D, S7E, S7F, S7G). Quantification showed that over 90% of the *in-situ* stiffened acini treated with integrin β1-blocking antibody or the FAK inhibitor remained non-invasive i.e. showed ∼9-fold reduction compared to the *in-situ*-stiffened control group, thus rescuing the phenotype completely (Figure 3F).

Taken together, these results show that *in-situ*-stiffening of normal mature mammary acini results in specific increase in activation of β1 integrins and signaling via activation of FAK which promote changes in the ECM composition and organization and drive the invasive phenotype.

### Mechanosensitive ion channels mediate *in-situ* stiffening induced invasion downstream of integrin activation

Multiple studies have demonstrated that MSCs such as TRP ion channels and Piezo1 are associated with ECM stiffness-driven cancer progression [25], [26]. Recent work also suggests significant crosstalk between MSCs and integrin-mediated focal adhesion signaling, which is known to transduce mechanical signals induced by increased ECM stiffness in various cell types [27], [28], [43]. As we found that integrin activation did not, on its own completely phenocopy the *in-situ* stiffened phenotype, we hypothesized that activation of MSCs might function as another effector in this process.

To test this, we inhibited the activity of MSCs using GsMTx4, a spider venom peptide which inhibits several MSCs including Piezo1 and TRP channels, in normal mammary acini during the stiffening process (Figure 4A). Single MECs were embedded in soft IPN gels, cultured to form mature acini over two weeks, and then *in-situ* stiffened with or without GsMTx4 for an additional two weeks. Confocal microscopy revealed that GsMTx4-treated acini maintained their circular morphology and hollow lumens, in contrast to the invasive phenotypes observed in untreated *in-situ* stiffened acini (Figures 4B, 4C, 4D, 4E, Figures S8A, S8B). Furthermore, GsMTx4 treatment reduced FN and increased LN levels, restoring their total and relative levels to those observed in normal acini in soft gels (Figure 4F, Figures S8C, S8D). Quantification confirmed that MSC inhibition significantly decreased the proportion of invasive acini, reducing the fraction of partially and completely invasive acini from ∼90% in untreated *in-situ* stiffened gels to ∼10% in GsMTx4-treated gels (Figure 4C). This was accompanied by a restoration of acini circularity and area to levels comparable to those in soft gels (Figures 4D, 4E, Figures S8A, S8B).

**Figure 4:**
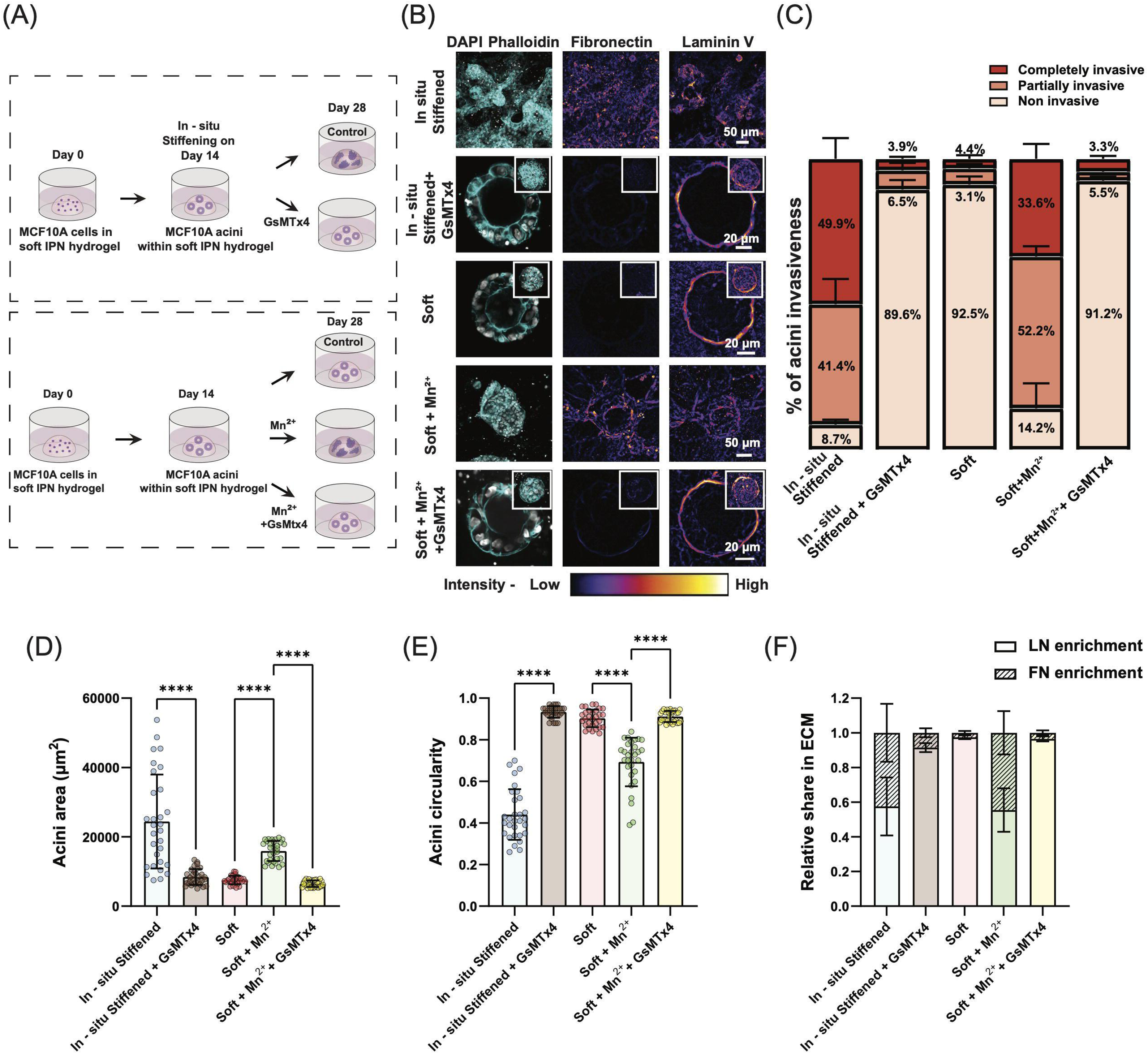
MSCs mediate in-situ-stiffening induced invasion downstream of integrin activation. (A) Schematic overview of the experimental overflow to test if MSCs are associated and crosstalk with integrins during the in-situ stiffening mediated invasion in normal mammary acini. Once the mature acini have formed in the soft IPN gels, some gels – (i) were in-situ stiffened which were then either cultured to serve as in-situ stiffened control or were incubated with media supplemented with GsMTx4, a MSCs inhibitor, (ii) served as soft control, (iii) were incubated with media supplemented with MnCl₂ (+/−) GsMTx4 to simultaneously block the activity of MSCs and activate integrins. (B) Confocal images showing nuclei (gray), F-actin (cyan), and FN and LN (fire LUT applied) in acini cultured under various experimental conditions. Z-stacked images are shown for acini in soft + Mn²⁺ and in-situ stiffened gels, while single Z-plane (middle plane image of the Z stack) images highlight lumen formation and retention in soft, soft + Mn²⁺ +GsMTx4, and in-situ stiffened + GsMTx4 conditions (Z-stacked images for the same acini in inset). Scale bars: 20 µm for soft, soft + Mn²⁺ +GsMTx4, and in-situ stiffened + GsMTx4; 50 µm for soft + Mn²⁺ and in-situ stiffened gel samples. (C-F) Quantification of (C) percentage of acini invasiveness, (D) acini area, (E) acini circularity, and (F) relative FN and LN enrichment in different experimental conditions (n = 30 acini from 3 different experiments, with 10 acini per experiment for area and circularity quantifications; n ≥ 30 acini from 3 experiments for invasiveness quantification; Kruskal-Wallis test, ****p < 0.0001).

To determine whether MSC activation was upstream or downstream of integrin and FAK activation, we then treated fully mature acini in soft gels with both MnCl_2_, which activates integrins independently of stiffness, and GsMTx4 (Figure 4A). Following the dual treatment, mammary acini showed no invasion maintaining their confined area, circularity, hollow lumen and relative levels of FN and LN, similar to normal mammary acini grown in soft gels (Figures 4B, 4C, 4D, 4E 4F, Figures S8A, S8B, S8C. S8D). This was in stark contrast to acini cultured in soft gels treated only with MnCl_2_ which exhibited the invasive phenotype and different LN and FN levels (Figures 4B, 4C, 4D, 4E, 4F, Figures S10). Quantification confirmed that while activating integrins with MnCl_2_ increased the number of partially invasive and completely invasive acini from ∼10% in soft gels to ∼85%, this was completely rescued by addition of GSMTx4 to ∼8% of invasive acini (Figure 4C). This rescue of invasive phenotype was through the same pathway of ECM regulation as addition of GSMTx4 to MnCl_2_ treated acini in soft gels significantly increased the levels of LN around the acini periphery in the ECM and reduced FN levels (Figures 4B, 4F). Taken together, these results confirm that activation of MSCs is an essential downstream target of *in-situ* stiffening and integrin activation that results in the invasive phenotype of normal mammary acini.

### Piezo1 drives invasion downstream of β1 Integrin–FAK signaling to regulate *in-situ* stiffening mediated changes in ECM composition and remodelling

Our findings discussed above demonstrate that *in-situ* stiffening induces the invasive phenotype in normal mammary acini through MSC activation downstream of integrin signaling. To identify the specific MSC mediating this process, we focused on Piezo1, a critical MSC known to interact with β1 integrin signaling pathways and regulate mechanotransduction [27], [29], [31]– [33], [44]. Piezo1 has also previously been implicated in ECM-driven cellular processes such as migration and invasion [32], [45], [46][29]–[33], making it a strong candidate for mediating stiffening-induced invasion in 3D mammary acini.

To investigate the role of Piezo1, we embedded single MECs in soft IPN gels and cultured them for 14 days to form normal mammary acini. We then *in-situ* stiffened the gels under five different conditions: (1) standard *in-situ* stiffened conditions for 14 days, (2) *in-situ* stiffened with β1 integrin blocking antibody (AIIB2), (3) *in-situ* stiffened with FAK inhibitor, (4) *in-situ* stiffened with β1 integrin blocking antibody plus the Piezo1 agonist Yoda1, and (5) *in-situ* stiffened with FAK inhibitor plus Yoda1 (Figure 5A). All samples were fixed, stained for LN and FN, and analyzed alongside normal mammary acini grown in soft gels for 28 days.

**Figure 5:**
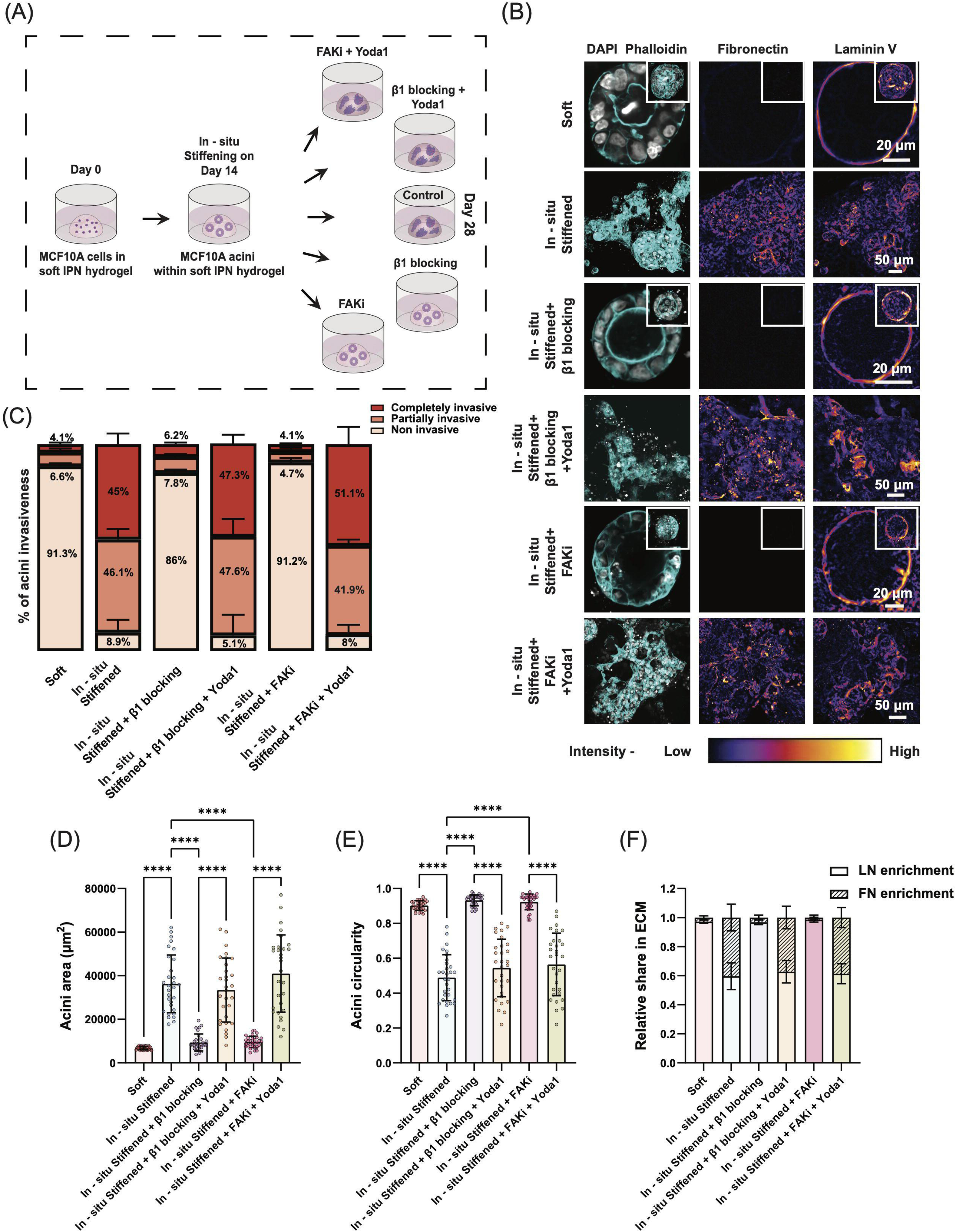
Piezo1 drives invasion downstream of β1 Integrin–FAK signaling to regulate in-situ stiffening mediated changes in ECM composition and remodelling. (A) Schematic overview of the experimental workflow to examine if Piezo1 MSC acts downstream to β1 Integrin–FAK Axis during the in-situ stiffening mediated mechanotransduction in normal mammary acini. After mature acini formation in soft IPN gels, gels were: (i) left untreated (soft control), (ii) in-situ stiffened and either left untreated (in-situ stiffened control), or treated with integrin β1 blocking antibody or FAKi (+/−) Yoda1, a Piezo1agonist. (B) Confocal images showing nuclei (gray), F-actin (cyan), and FN and LN (fire LUT applied) in acini cultured under various experimental conditions. Z-stacked images are shown for acini in in-situ stiffened, in-situ stiffened + β1 blocking + Yoda1, and in-situ stiffened + FAKi + Yoda1 conditions, while single Z-plane images are shown to highlight lumen formation and retention in soft, in-situ stiffened + β1 blocking, and in-situ stiffened + FAKi conditions (Z-stacked image for the same acini in inset). Scale bars: 20 µm for soft, in-situ stiffened + β1 blocking, and in-situ stiffened + FAKi; 50 µm for in-situ stiffened, in-situ stiffened + β1 blocking + Yoda1, and in-situ stiffened + FAKi + Yoda. (C-F) Quantification of (C) percentage of acini invasiveness, (D) acini area, (E) acini circularity, and (F) relative FN and LN enrichment in different experimental conditions (n = 30 acini from 3 different experiments, with 10 acini per experiment for area, circularity and FN and LN enrichment quantifications; n ≥ 30 acini from 3 experiments for invasiveness quantification; Kruskal-Wallis test, ****p < 0.0001).

Confocal microscopy revealed that as shown above, *in-situ* stiffening of normal mammary acini resulted in invasive phenotypes which were rescued by blocking activation of β1 integrin or inhibition of FAK (Figure 5B). However, we observed that this blocking effect was lost in the presence of Yoda1, both in terms of morphology of the acini as well as levels of LN and FN along the invading fronts in the ECM (Figure 5B). Quantification of acini invasiveness showed that while inhibiting β1 integrin-FAK signaling reduced partially and completely invasive acini by ∼ 6-7-fold, this effect was lost in the presence of Yoda1 which resulted in complete restoration of invasive acini to levels seen in *in-situ* stiffened conditions as well as in overall acini size and circularity (Figures 5C, 5D, 5E, Figures S9). Additionally, quantification of LN and FN showed that activation of Piezo1 in *in-situ* stiffened acini in the presence of β1-FAK inhibition was sufficient to down-regulate LN levels and upregulate FN levels in the ECM which correlated with the increased invasiveness observed under these conditions (Figures 5F, Figures S9). Taken together, these results show that Piezo1 is a key MSC that acts downstream of *in-situ* stiffening mediated β1 integrin and FAK signaling and regulates ECM composition to drive the invasive phenotypes in mammary acini.

## Discussion

Here we show that *in-situ* stiffening of normal mammary acini triggers distinct invasive phenotypes through dynamic ECM remodeling. This process begins with stiffening-induced priming of the acinar environment, characterized by downregulation or degradation of the BM and increased stromal ECM production around the acini structure. This is followed by a second phase marked by increase in cellular proliferation and migration that coincides with upregulated BM production and further stromal ECM remodeling. Mechanistically, we find that *in-situ* stiffening induced ECM remodeling and invasion is triggered by increased activation of FN-binding β1 integrins and its downstream kinase FAK. Critically, we identify that the MSC Piezo1 is the downstream effector of this process, activation of which in the presence of the stiffened ECM is necessary and sufficient to drive invasion.

Interestingly, fully formed acini subjected to *in-situ* stiffening in our study exhibit invasive behaviors distinct from the invasive phenotypes observed when single cells are grown on stiff environments. Unlike the complete absence of BM observed in invasive acini grown from single MECs in stiff gels, here we find that *in-situ* stiffened acini have high levels of BM that is remodeled and localized around the invasive structures. These findings align with previous studies showing that abnormal ECM mechanics can compromise the BM barrier through protease degradation [47] or mechanical forces from transformed cells [10], [48]. The other difference we observe between the invasive phenotype in stiff vs *in-situ* stiffened acini is the heterogeneity of phenotypes. Here, we find that *in-situ* stiffening results in some acini being partially invasive, some completely invasive and some not responding to changes in stiffness at all which contrasts with acini grown from single MECs on stiff gels where most acini are completely invasive. We currently don’t understand mechanistic basis of this heterogeneity, although it may be that the ability of normal acini to respond to stiffening is determined by the size of an acini or its cellular organization and state within the acini. It is also tempting to speculate here that *in-situ* stiffening of acini may be more representative of tumor growth and invasion at the primary site while single cells growing and expanding in stiff gels may reflect metastasis at secondary sites which often originate from single or a group of metastasized cells.

By following the *in-situ* stiffening mediated invasion process over 14 days, we identify two key features about the dynamics of the process. The first is the dramatic changes in BM and stromal ECM levels immediately post stiffening, where we find FN levels to immediately increase while LN levels first decrease significantly. We propose calling this the pre-invasion ECM priming phase. This is followed by the second phase which is the onset of cell proliferation and invasion which initiates ∼6 days post stiffening, coinciding with upregulation of BM and significant remodeling of the stromal ECM. This shift in BM levels implies a functional transition in LN, from maintaining epithelial polarity to supporting migration and tumorigenesis, with the mechanistic basis of this biphasic behavior remain to be clearly identified. The dynamic changes in the BM are coupled to rapid increase in FN secretion upon stiffening. While we experimentally interpret these 2 phases to be distinct, it is more likely that the phases overlap, with cellular changes occurring in parallel to changes in the ECM environment. We can also assume that this initial process is independent of FN-binding integrins such as β1 since we find negligible levels of FN in the normal mammary acini. This leads us to suggest two possible mechanisms, one that is via BM-dependent mechanotransduction pathways or the other that is initially integrin independent. Since several studies have now shown that LN binding integrins are mechanosensitive and can respond to changes in the mechanics of the ECM [49], the BM-dependent pathway could be through stiffness-mediated activation of integrins such as α6β4 that eventually leads to upregulation of FN secretion. The other possible mechanism is through an upstream MSC since studies have previously shown that MSCs such as Piezo 1 could also result in upregulation of FN secretion thus potentially presenting a pathway that is Integrin-independent [44].

Lastly, our results specifically place Piezo 1 activation downstream of β1 integrin-FAK pathway and as the critical regulator of *in-situ* stiffening mediated invasion. Activation of Piezo 1 has been implicated in several cancers [25]–[27], [29], [32] but mechanisms by which Piezo 1 can be activated downstream of integrins remains unknown. Studies have now shown significant cross talk between integrin and Piezo 1 pathways [29]–[33] such as through local changes in membrane tension in proximity of focal adhesions [29]. It is also likely that Piezo 1 and integrins interact with each other via more complex feedback mechanisms that regulate and amplify each other and this needs to be probed in greater detail.

In summary, our *in-situ* stiffening system and the mechanisms discovered here following its application, have implications not only in the context of cancer and the onset of invasion but also in the context of development and the morphological changes that occur in the sculpting of complex tissues. Importantly, our findings highlight potential therapeutic strategies targeting mechanotransduction pathways involved in stiffening-induced cellular responses. For instance, Piezo1 inhibitors could be explored to mitigate the invasive behavior of tumorigenic cells by reducing mechanosensitive calcium influx, thereby limiting their ability to breach the BM. Similarly, anti-β1 integrin antibodies may offer a viable approach to disrupt integrin-mediated mechanotransduction, which is critical for cell-ECM adhesion and signaling, ultimately reducing tumor cell invasion and metastasis. These insights underscore the relevance of targeting cellular mechanosensing mechanisms in both cancer therapy and tissue engineering applications.

## Methods

### Cell culture

MCF10A and MCF10A DCIS.com cells, a gift from Dr. Guillaume Jacquemet (Åbo Akademi University), were cultured in Dulbecco’s modified Eagle’s medium/Nutrient Mixture F-12 (DMEM/F12) medium (Thermo Fisher) supplemented with 5% horse serum (Thermo Fisher), 20Lng/ml EGF (Miltenyi Biotech), 0.5 µg/ml hydrocortisone (Sigma), 100Lng/Lml cholera toxin (Sigma), 10 µg/ml insulin (Sigma) and 1% penicillin/streptomycin (Thermo Fisher). Cells were routinely passaged every 3–4Ldays with TrypLE™ Express Enzyme (Thermo Fisher) and cultured in a standard humidified incubator at 37□°C and 5 % CO_2_.

### Alginate preparation

1% w/v unmodified sodium alginate (Pronova UP MVG, Novamatrix), rich in guluronic acid and with high molecular weight, was dissolved in ultrapure water, sterile filtered and lyophilized. Subsequently, reconstituted alginate stock solution was prepared by dissolving the lyophilized alginate in serum free DMEM/F12 media overnight to bring the total alginate concentration to 2.5% w/v.

### IPN gel preparation

To prepare IPNs with final concentrations of 5 mg/mL alginate and 4.5 mg/mL BME (Cultrex Basement Membrane Extract, Type 2, Pathclear, Biotechne R&D Systems), prechilled reconstituted alginate was mixed with BME and serum-free DMEM/F12 media in a 1.5 mL microcentrifuge tube on ice. The mixture was transferred to a 1 mL Luer Lock syringe (Fisherbrand™). Separately, serum free DMEM/F12 media with or without 20 mM CaSO_4_ was loaded into a second 1 mL Luer Lock syringe for preparing stiff and soft IPNs, respectively. The two syringes were coupled using a female–female Luer-Lock connector (Masterflex®, VWR), mixed rapidly, and immediately deposited into appropriate dishes for AFM and PTM measurements. The solutions were allowed to gelate for 1 hour in an incubator at 37 °C to form stiff and soft IPN gels. If required, soft gels were *in-situ* stiffened by incubating them in a 20 mM CaCl_2_ solution for 30 minutes. The *in-situ* stiffened gels were washed thrice with phosphate buffered saline (PBS).

### Mechanical testing

*AFM Measurements* - AFM force spectroscopy experiments were performed with a NanoWizard 4 (JPK, Bruker) mounted on a Nikon Ti-2 inverted microscope with a x20 objective (Nikon ELWD S Plan Fluor, NA=0.6). Measurements were performed with rectangular cantilevers with a colloidal probe of radius R = 2.25 µm (NanoAndMore, CP-qp-CONT-Au-A). The cantilevers have a nominal resonance frequency of 30 kHz and nominal spring constant 0.1 N/m. The gels were indented with a setpoint value of 0.2-1.0 nN with a speed of 1 µm/sec and sampling rate of 1000 Hz. Measurements were performed in PBS with no magnesium and no calcium.

The analysis of the force-distance curves is based on a previous study [50] and uses the following relation between the measured force *F* and indentation δ for a parabolic indenter geometry

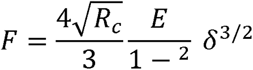

where *E*, □, and *R_c_* are the Young’s modulus of the sample, the Poisson’s ratio of the sample, and the effective radius of the indenter tip curvature, respectively. Poisson’s ratio is assumed to be 0.5.

*PTM measurements –* PTM was used to map the dynamics of soft IPN gel stiffening upon crosslinking. PTM involves tracking the thermally driven Brownian motion of probe particles over time, using their trajectories to extract the viscoelastic properties of the surrounding material.

To image the probe particles, a widefield microscope equipped with a x60 objective lens (Olympus, numerical aperture 1.1, working distance 1.5 mm) and an sCMOS camera (Hamamatsu, ORCA Flash 4.0) was used. Soft IPN gels were prepared as previously described, with the addition of 6 µm diameter polystyrene beads to the gel solution to serve as probe particles after gelation. The prepared soft IPN gel with embedded probe particles was deposited into Ibidi μ-Slide 4-well glass-bottom chambered coverslips.

Measurements were recorded over a 20-hour period, after which the soft gels were *in-situ* stiffened by the addition of 20 mM CaCl2 at the 22-hour time point. Following a 30-minute incubation in the CaCl2 solution, the gels were rinsed three times with sterile MilliQ water.

For each rheological measurement, an individual probe particle was selected, and a small region of interest (20 × 20 µm) was imaged at a frame rate of approximately 300 frames per second. A total of 300,000 frames were collected per measurement. A schematic of the imaging system is shown in Figure 2, which also includes an example of a small region of interest containing a probe particle. The Brownian motion of the probe particle was tracked using a center-of-mass algorithm (Figure 2). To extract the low frequency elastic modulus of the material local to the probe the analysis steps detailed in previous PTM studies [37], [38]. The mean squared displacement, MSD, of the probe’s motion is analysed, and related to the time-dependent compliance of the material local to the probe, J(t), as follows,

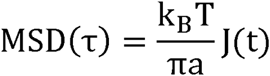

where τ is the lag-time, k_B_ - Boltzmann constant, T - temperature, and a - the radius of the probe. The Fourier transform of the time-dependent compliance, J(ω), is used to determine the complex modulus, G*(ω) of the material, an example of which is shown in Figure 2A,

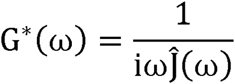

where the real, G’(ω), and imaginary, G’’(ω), components of the complex modulus relate to the elastic and viscous response of the material respectively. In Figures 2A, we compare the elastic properties of different samples by plotting the low frequency elastic modulus (G_0_’), in other words the low frequency, long time, plateau in G’(ω).

### Acini formation in IPN gels

To form acini, soft and stiff IPN gels were prepared as described above, with the addition of a single-cell suspension of MCF10A cells at a density of 35,000 cells/mL to the mixture within the first syringe. After mixing the contents of both syringes, the IPN gel mixture with MCF10A MECs was transferred to a 1.5 mL microcentrifuge tube. Subsequently, 30 µL of the soft IPN gel mixture were pipetted into each well of a 24-well plate precoated with BME (1:5 dilution in serum-free DMEM/F12 media) to form dome-like drops. The plate was incubated at 37 °C for at least one hour to allow the formation of soft IPN gels. Once the gels were set, acini culture media (DMEM/F12 media supplemented with 2% horse serum, 5 ng/mL EGF, 0.5 µg/mL hydrocortisone, 100 ng/mL cholera toxin, 10 µg/mL insulin, and 1% penicillin/streptomycin) was carefully added to the sides of the wells to prevent gel detachment. The media was replaced every 3 to 4 days throughout the culture period. Once acini had formed within the soft IPN gels by day 14, a subset of these gels was subjected to stiffening or treated with specific pharmacological agents to investigate the role of integrin or Piezo1 activation and exogenous FN on normal acini. For stiffening, the acini culture media was replaced with a 20 mM CaCl_2_ solution, and the soft IPN gels were incubated in this solution for 30 minutes at 37 °C. After incubation, the CaCl_2_ solution was removed, and the gels were washed three times with PBS before fresh acini culture media, with pharmacological agents that promote or inhibit invasion in *in-situ* stiffened conditions, if required, was added. The acini were then cultured for an additional 14 days in the *in-situ* stiffened gels.

To form acini in soft IPN gels pre-mixed with FN, the same procedure as described above was followed. However, a FN solution was added to the contents of the first syringe, achieving a final concentration of 20 µg/mL in the soft IPN gel mixture.

### Acini formation in BME gels

A single-cell suspension of MCF10A MECs was prepared by mixing the cells with 4.5% w/v BME (diluted in serum-free DMEM/F12 medium) in a 15 mL centrifuge tube on ice to achieve a final cell density of 35,000 cells/mL. The mixture was pipetted as dome-shaped drops into each well of a 24-well plate. The cells were cultured for 14 days to allow acini formation. On day 14, a subset of the BME gels containing fully formed acini was incubated with a 20 mM CaCl_2_ solution to investigate whether calcium induces invasive properties in normal acini within a soft microenvironment. After a 30-minute incubation at 37 °C, the BME gels were washed thrice with PBS, and fresh acini culture media was added. The acini within the BME gels were cultured for an additional 14 days.

### Pharmacological inhibition

Pharmacological agents were added to the acini culture media on day 14 to treat normal acini in soft gels or *in-situ* stiffened IPN gels. The concentrations of pharmacological agents for treating normal acini in soft and *in-situ* stiffened IPN gels are as follows: Soft IPN gels - 0.5 mM MnCl_2_ (Sigma) for integrin activation, 10 µM Yoda1 (Tocris Bioscience) for Piezo1 activation, 20 µg/mL exogenous fibronectin (Sigma) for assessing the effects of exogenous FN Stiff IPN gels – 5 µg/mL Anti-Integrin β1 clone AIIB2 (Sigma) for β1 integrin inhibition, 5µM PF-573228 (Sigma) for FAK inhibition and 5 µM GsMtx4 (MedChem Express) for Piezo1 inhibition.

### Immunofluorescence staining and imaging

To fix IPN gel samples embedded with acini for immunofluorescence staining, 2% paraformaldehyde (Thermo Fisher) in PBS supplemented with 0.1% glutaraldehyde (Sigma) was added to the gels and incubated for 30 minutes at room temperature. The gels were washed three times with PBS and then permeabilized using 0.15% Triton X-100 (Fisher Scientific) for 25 minutes. Afterward, they were washed again three times with PBS and incubated overnight in a blocking buffer consisting of 3% BSA solution in PBS.

The gels were subsequently incubated with primary antibodies (1:400), diluted in 1% BSA and 0.3% Triton X-100 in PBS, overnight at 4 °C. After washing three times with PBS, the gels were incubated in a solution containing fluorescently conjugated secondary antibodies, Alexa Fluor 488–phalloidin (1:400, Thermo Fisher), and Hoechst 33342 solution (1:4000) for 90 minutes at room temperature, followed by three PBS washes. The gels were mounted with ProLong™ Glass Antifade Mountant and sandwiched between two round coverslips.

Images of the gel samples on coverslips were acquired using a Nikon Confocal A1RHD microscope equipped with 405-, 488-, 561-, and 640-nm laser lines and GaAsP PMTs, using x10 and x20 objectives with numerical apertures of 0.45 and 0.8, respectively.

Primary antibodies used included anti-fibronectin (from rabbit, Sigma, 1:400), anti-laminin-5 (γ2 chain, clone D4B5, from mouse, Sigma, 1:400), anti-β1 integrin (from mouse, Sigma, 1:400), and anti-β4 integrin (clone 439-9B, from rat, Thermo Fisher, 1:400). Secondary antibodies used were Alexa Fluor 568 goat anti-mouse IgG (Thermo Fisher, 1:400), Alexa Fluor 568 goat anti-rabbit IgG (Thermo Fisher, 1:400), Alexa Fluor 647 goat anti-mouse IgG (Thermo Fisher, 1:400), and Alexa Fluor 647 goat anti-rabbit IgG (Thermo Fisher, 1:400).

### Image analysis and quantification of acini phenotypical parameters

All image analyses, including the quantification of acini area, acini circularity, and mean intensities of FN and LN expressed by acini, were performed in ImageJ. x10 and x20 image sequences acquired with the confocal microscope were converted into Z-projection (maximum intensity projection) images and background-subtracted.

Acini images from at least three biological replicates per experimental condition were used for quantification. Acini were manually categorized as partially invasive, completely invasive, normal, or non-invasive based on descriptions provided (see results). Acini area and circularity were measured by manually tracing the outline of the F-actin/Phalloidin channel of Z-projected acini images in ImageJ.

The mean intensities of FN, LN, β1 integrin, and β4 integrin were calculated by measuring the mean pixel intensity of their respective staining within the same manually drawn regions of interest used for area and circularity measurements. Normalized FN and LN enrichment were calculated using the formula:

Normalized FN or LN Enrichment = Mean Intensity of FN or LN of an acinus/ Sum of Mean Intensities of FN and LN of the same acinus

A Fire LUT was applied to Z-stacked, background-subtracted FN and LN staining images to visualize ECM remodeling in representative acini across all figures.

### Statistical analysis

All data were analyzed using GraphPad Prism version 10 (GraphPad Software, Boston, Massachusetts USA, www.graphpad.com). The specific number of acini per biological replicate for each experimental condition is indicated as *n* in the respective figure legends. Non-normally distributed data were analyzed using a Mann-Whitney U test. Scatter plots with bars, stacked bar plots, and dot plots were used for data display, with horizontal lines representing the mean.

Statistical tests used for each experiment are displayed in the corresponding figure legends. Asterisks indicate statistical significance, with the following *p*-value ranges: **** P < 0.0001, *** P < 0.001, ** P < 0.01, * P < 0.05, ns, not significant.

## Acknowledgements

We thank Drs. Martijn Gloerich, Guillaume Jacquemet, Darcy Wagner, Sebastian Wrighton, Berit Olofsson and Sebastian Wasserstrom as well as all the members of laboratory of cell and molecular mechanobiology (LCMM) for their advice and support. Lund University Bioimaging Center (LBIC) at Lund University is gratefully acknowledged for providing imaging resources. This research was funded by the Knut and Alice Wallenberg Foundation (V.S.) via the Wallenberg Centre for Molecular Medicine, Lund; Cancerfonden (VS, 19 0445 Pj and 22 2398 Pj Projekt grant), Fru and Berta Kamprad Foundation (VS, FBKS-2023-31), Crafoord Foundation (VS), The Royal Physiographic Society of Lund (KS), Novo Nordisk Foundation (ACS, NNF21OC0071368) and EPSRC grants EP/R035067/1, EP/R035563/1, and EP/R035156/1(AJW).

