## Supplementary figures and images for "β1 Integrin–FAK–Piezo1 signalling axis drives *in-situ* stiffening mediated ECM remodelling and invasion of 3D breast epithelium"

### Supplemental Figure 1

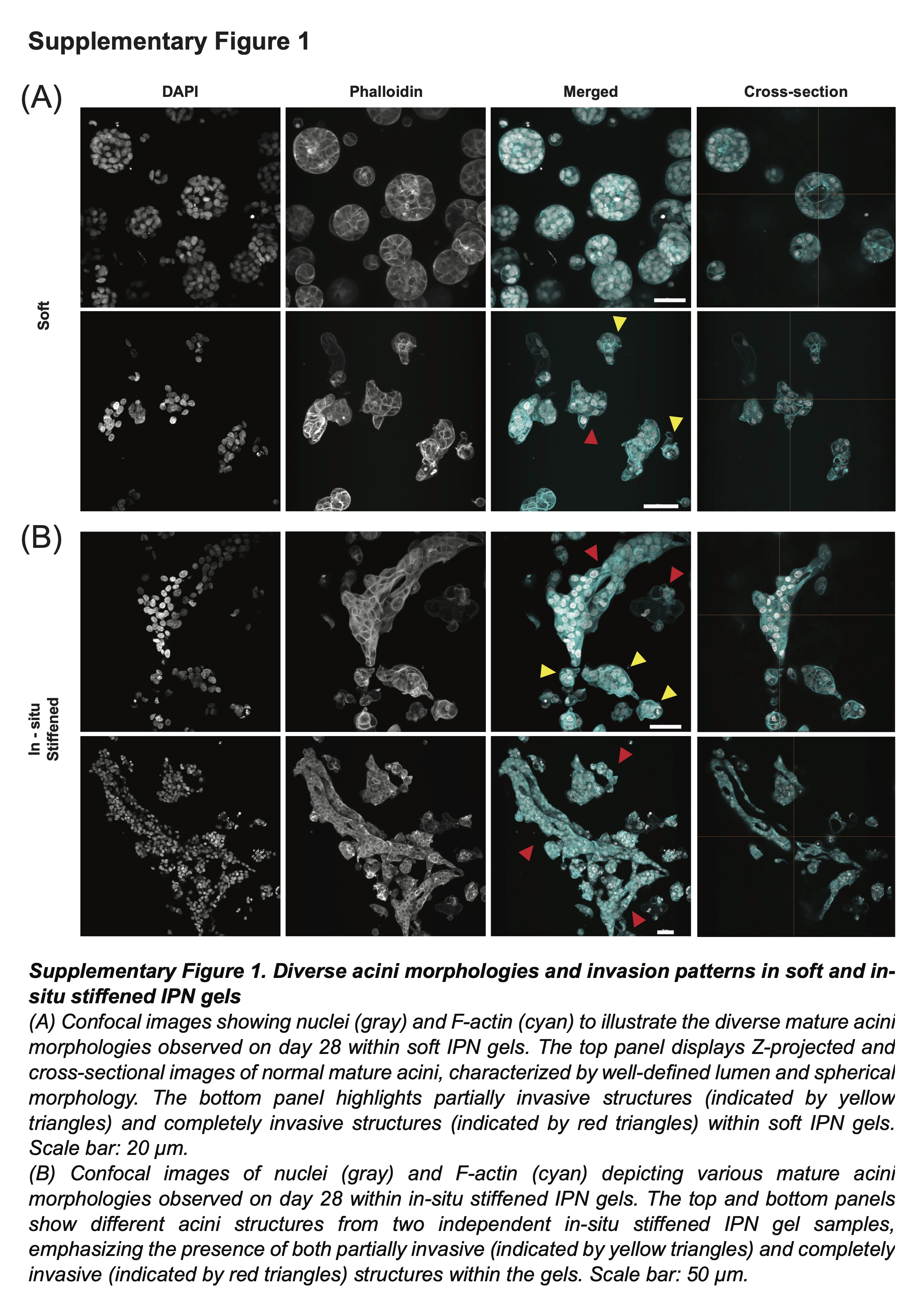

### Supplemental Figure 2

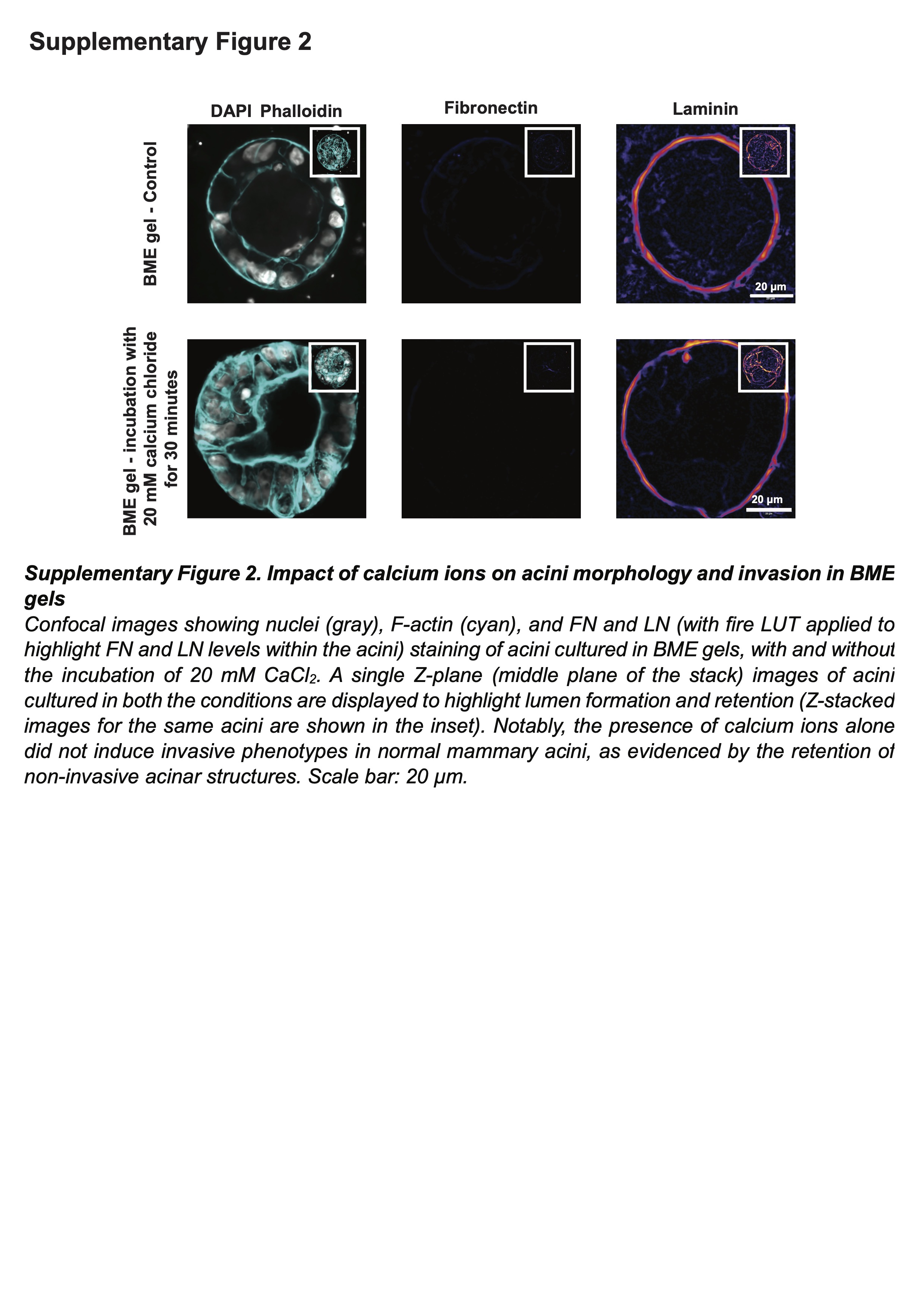

### Supplemental Figure 3

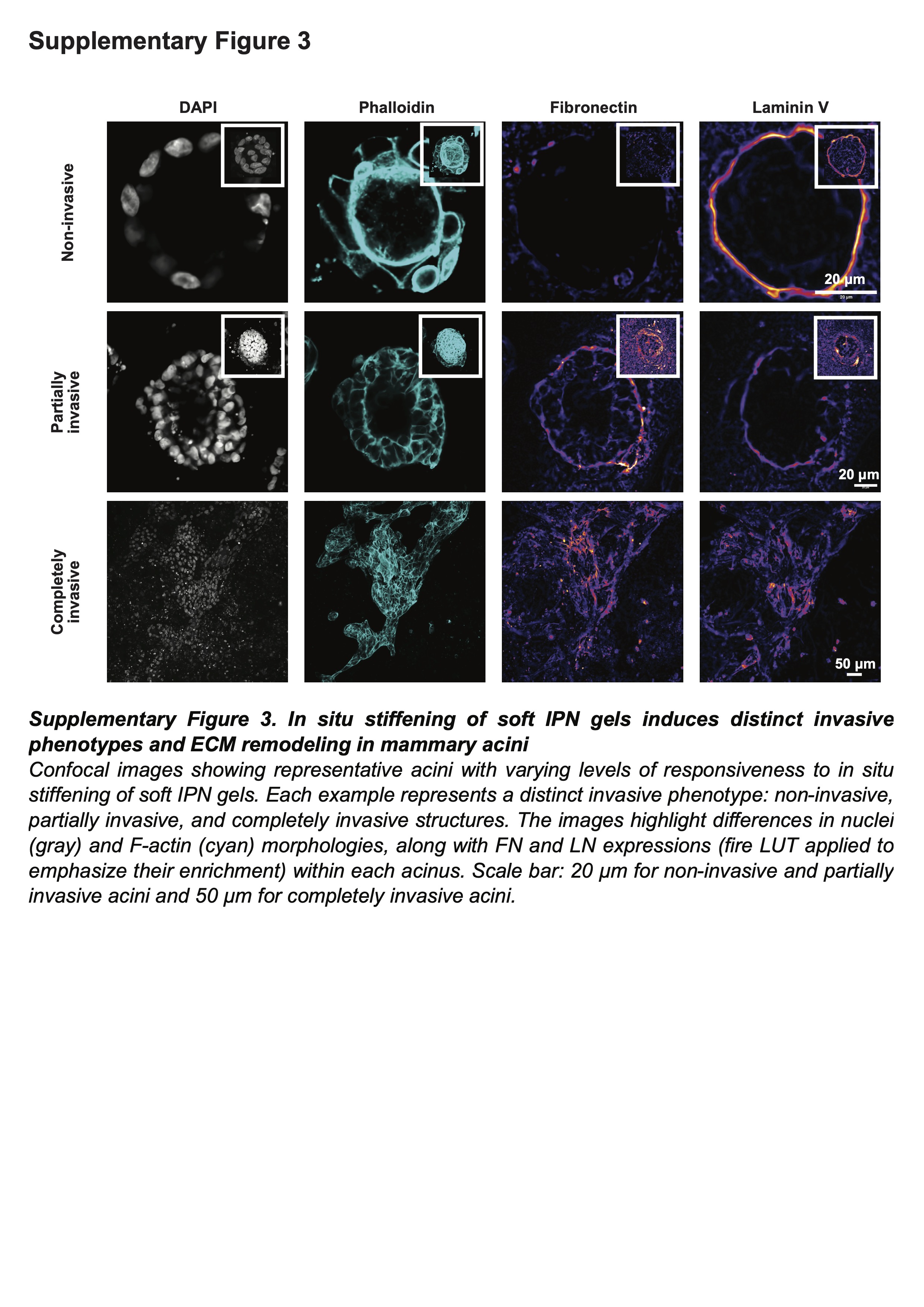

### Supplemental Figure 4

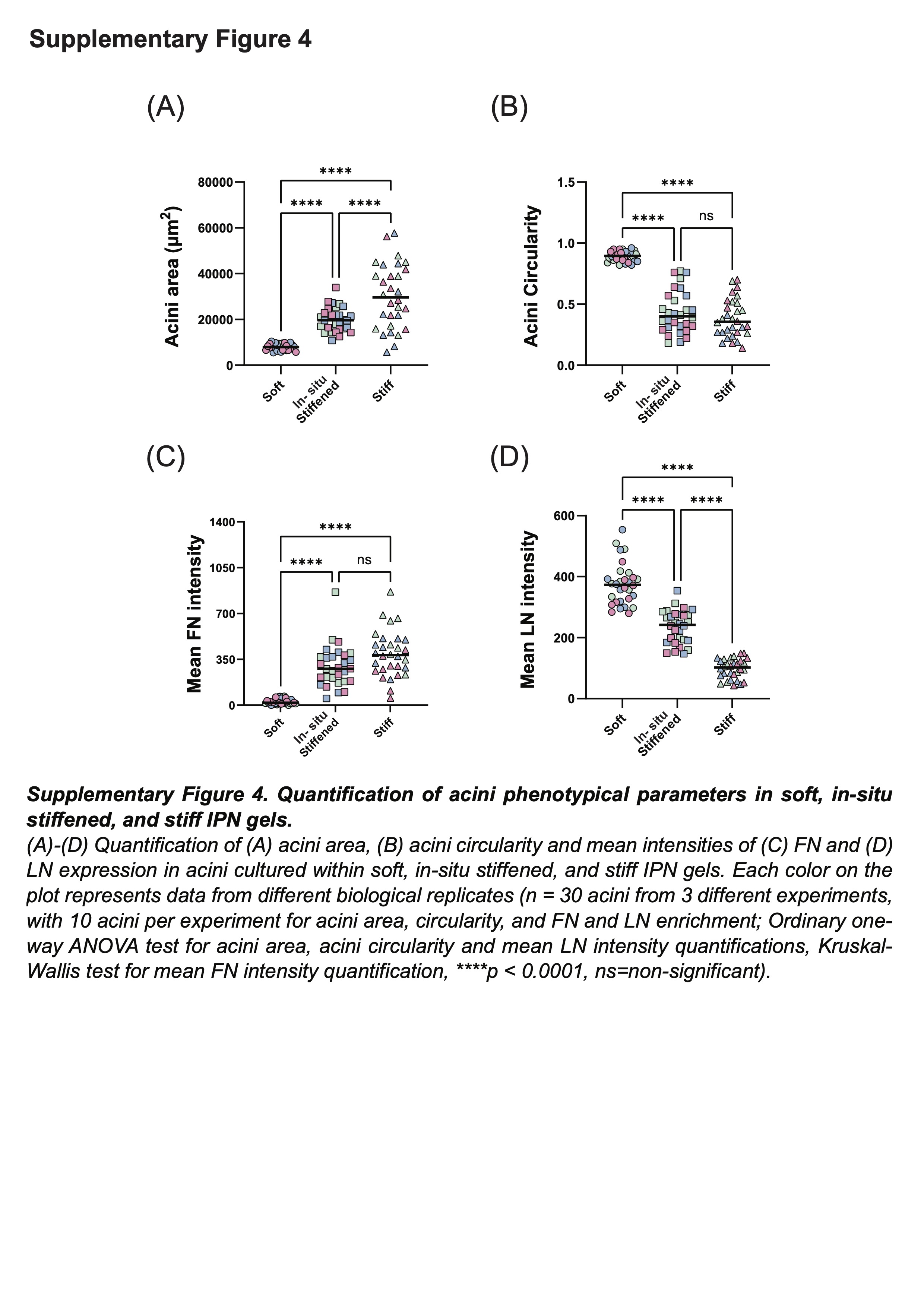

### Supplemental Figure 5

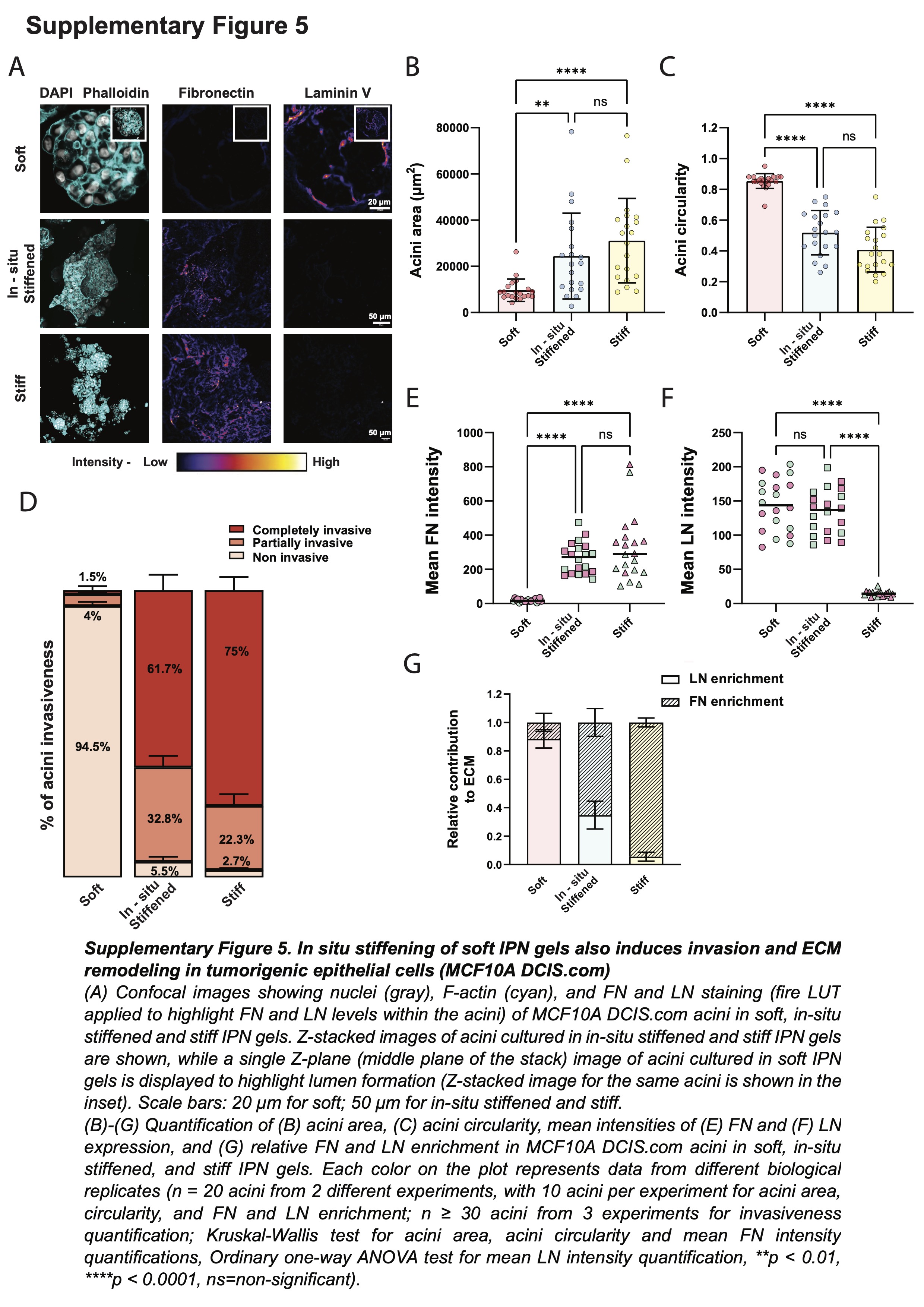

### Supplemental Figure 6

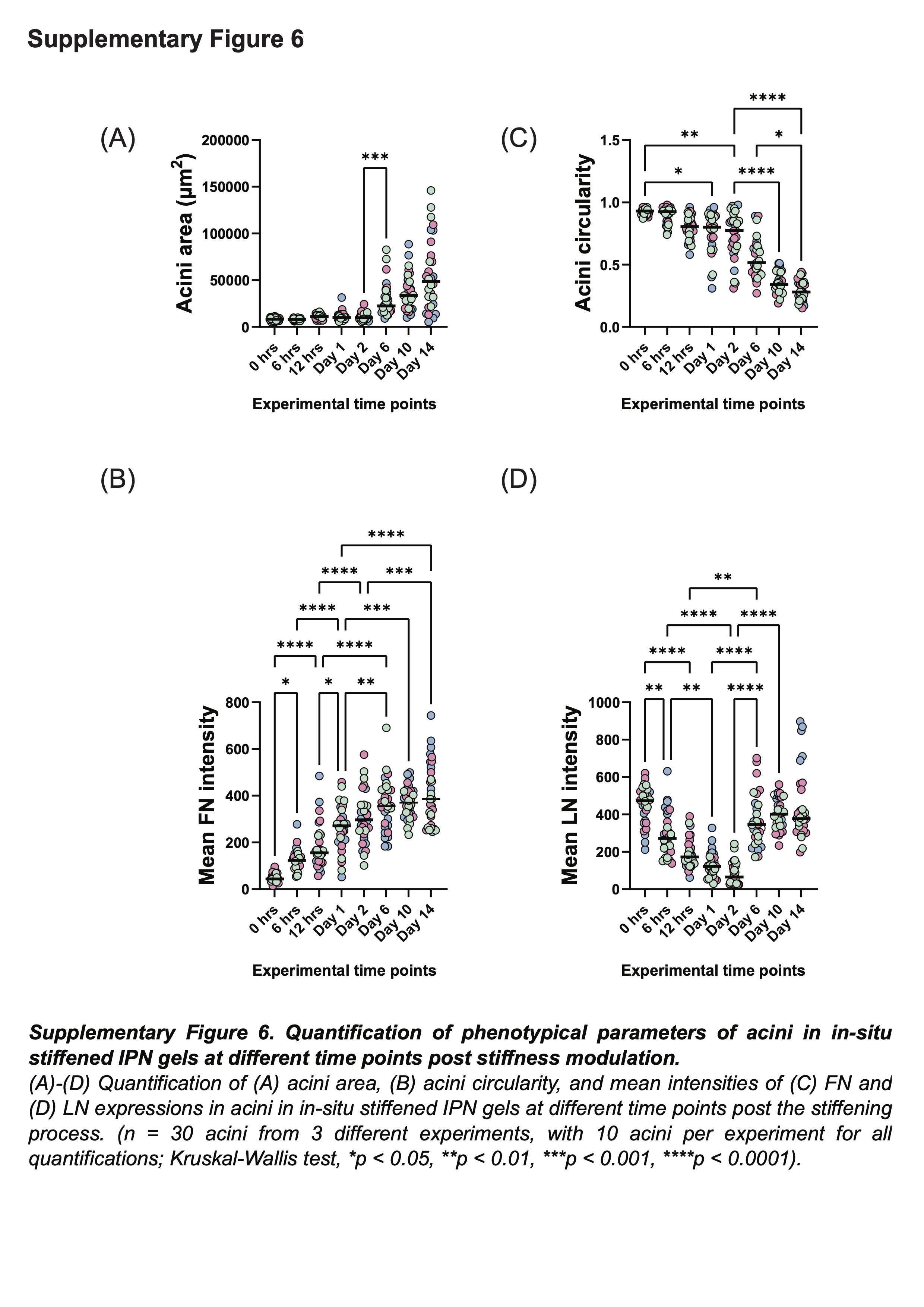

### Supplemental Figure 7

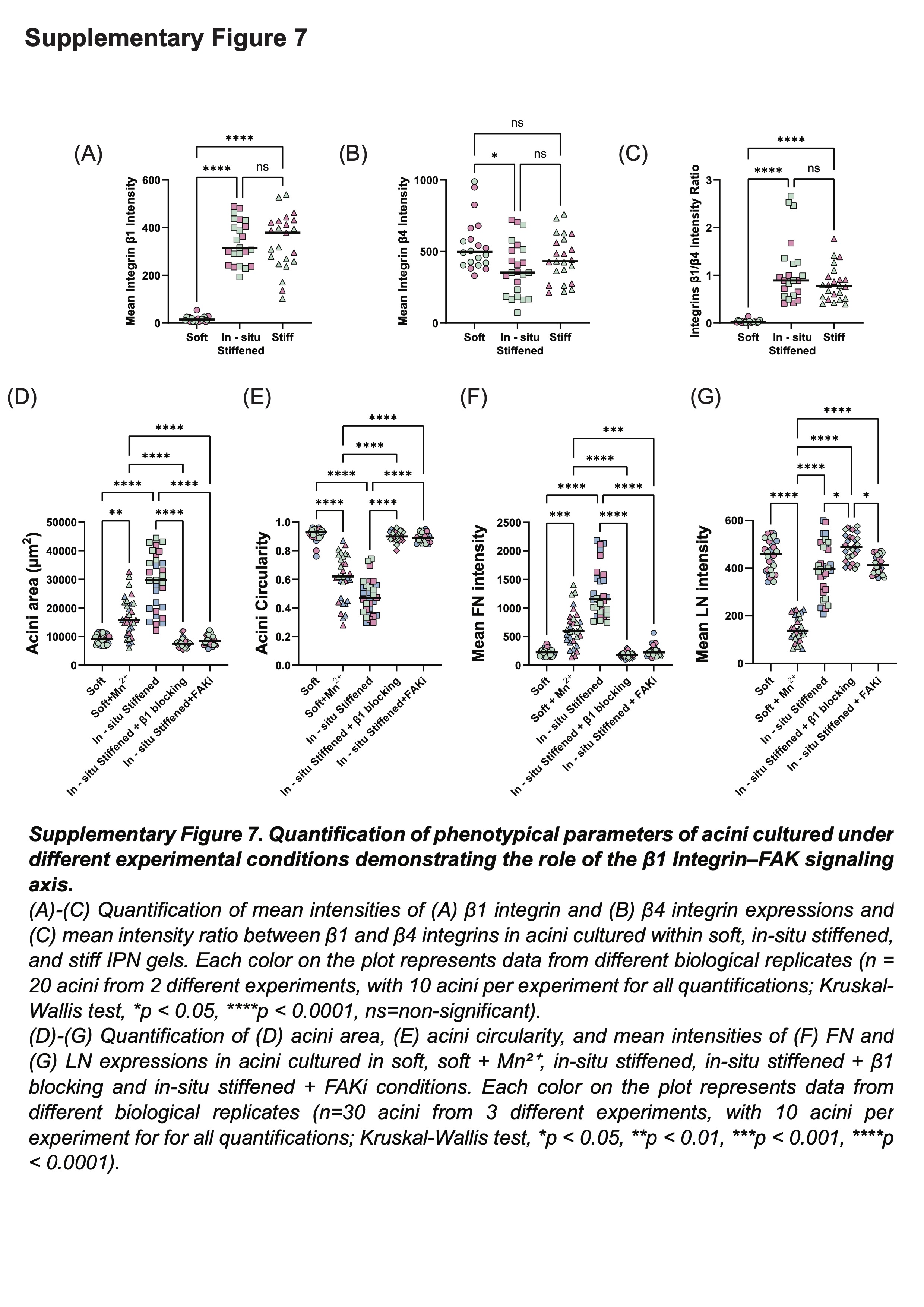

### Supplemental Figure 8

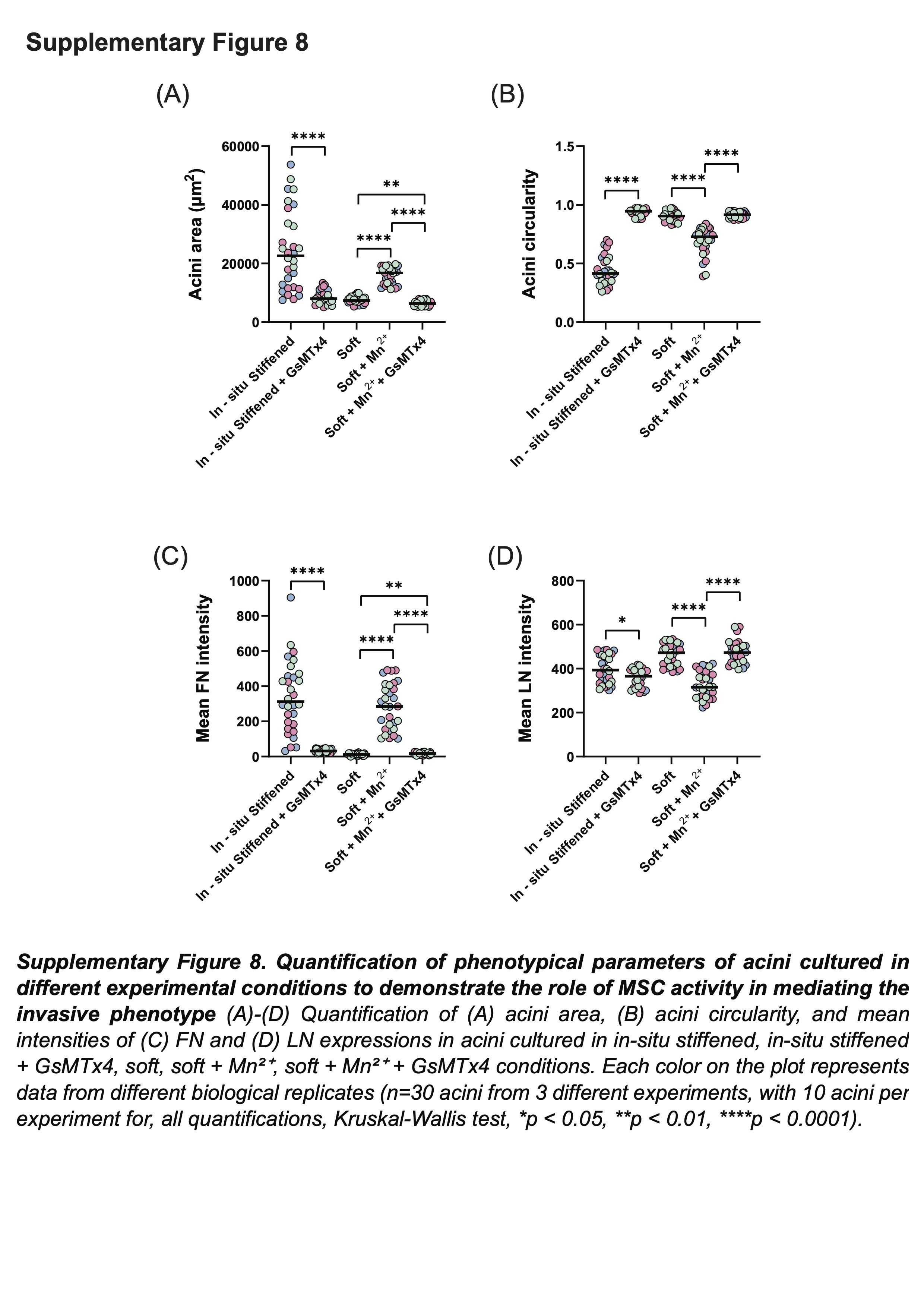

### Supplemental Figure 9

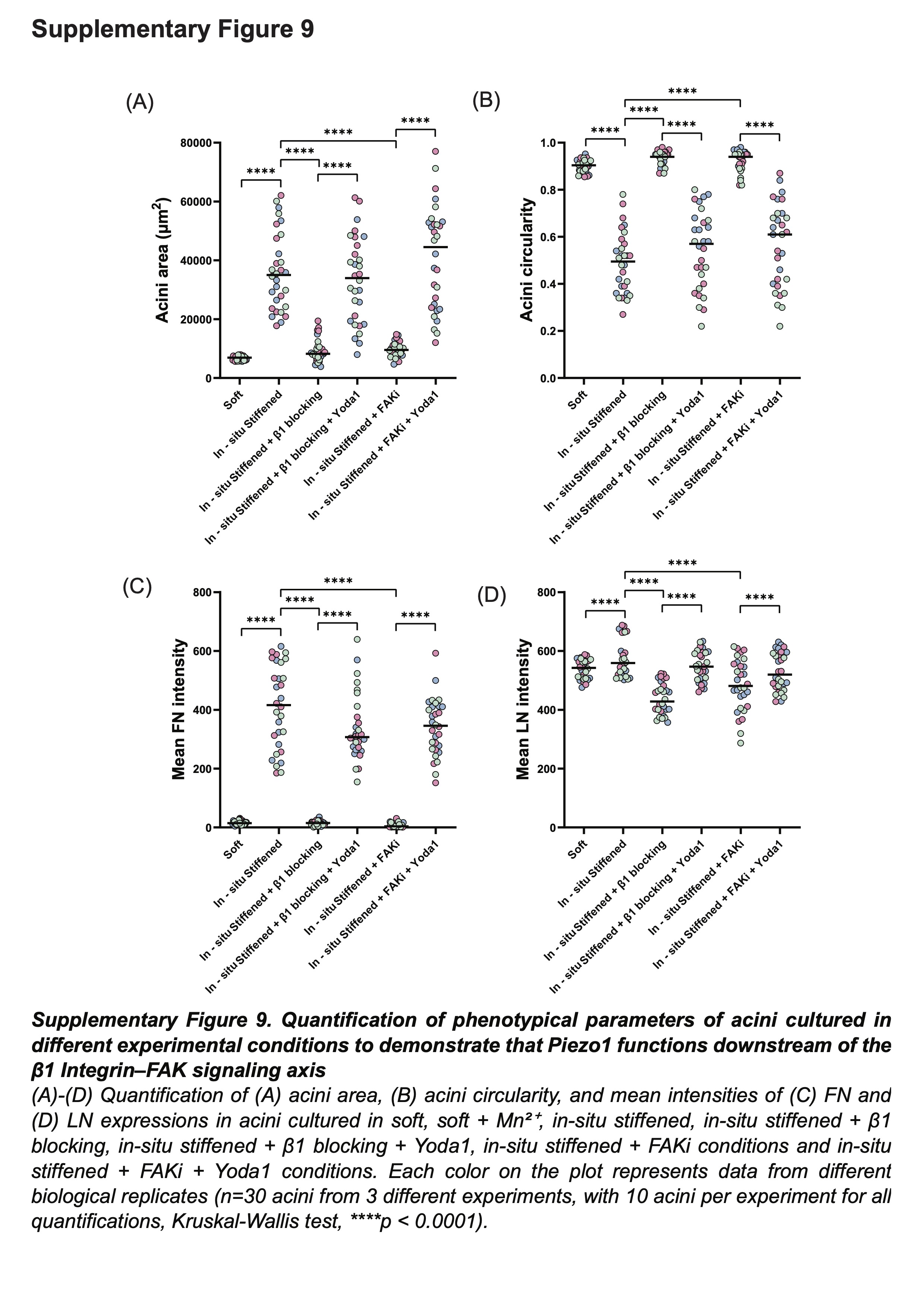
